# Learning interpretable representation for context-specific transcription regulatory networks using a foundation model

**DOI:** 10.1101/2025.03.29.646077

**Authors:** Zhaowei Yu, Dongxu Yang, Qianqian Chen, Yuxuan Zhang, Zhanhao Li, Yucheng Wang, Chenfei Wang, Yong Zhang

**Author notes:** These authors contributed equally to this work (Zhaowei Yu, Dongxu Yang, Qianqian Chen, Yuxuan Zhang). Correspondence: Yong Zhang Ph.D., 1239 Siping Road, Shanghai 200092, China.

## Abstract

Gene expression is shaped by transcription regulatory networks (TRNs), where transcription regulators interact within regulatory elements in a context-specific manner. Despite significant efforts, understanding the intricate interactions of transcription regulators across different genomic regions and cell types remains a major challenge, largely due to data sparsity. Here, we introduce ChromBERT, a foundation model pre-trained on large-scale human ChIP-seq datasets. ChromBERT effectively captures the interaction syntax of approximately one thousand transcription regulators across diverse genomic contexts, generating interpretable representations of context-specific TRNs and their constituent regulators. Fine-tuned for various transcription regulation tasks, ChromBERT demonstrates superior performance in imputing previously unseen cistromes and modeling regulatory effects and dynamics in specific cell types. Notably, ChromBERT adapts its TRN representations for cell-type-specific tasks, providing deep insights into the roles of transcription regulators underlying observed regulatory effects or dynamics without requiring cell-type-specific genomic data for each regulator. By overcoming the limitations of sparse transcription regulator data, ChromBERT significantly enhances our ability to model, interpret, and predict transcriptional regulation across diverse biological contexts.

## Introduction

Gene expression patterns are shaped by the intricate interactions among transcription regulators, regulatory elements, and target genes, collectively forming transcription regulatory networks (TRNs)^1–4^. A central goal in biology is to understand how transcription regulators interact within specific genomic contexts, how distinct regulatory architectures are formed in different genomic regions, and how these interactions vary across diverse cell types. These interactions underlie cell-type-specific transcriptional programs, which are critical for maintaining cellular identity and function. Landmark projects like the Encyclopedia of DNA Elements (ENCODE) have provided invaluable insights into transcription regulatory landscapes by generating extensive data on transcription factor binding and chromatin states^2,4^. However, these efforts have primarily focused on a limited number of well-studied cell types. Consequently, the context-dependent interactions of transcription regulators and their hierarchical roles in modulating gene expression remain poorly understood in most biological settings, particularly in dynamic processes such as cellular transitions and disease progression.

Pre-trained foundation models have achieved remarkable success in genomics studies by leveraging large-scale heterogeneous datasets to learn fundamental knowledge that can be fine-tuned for a wide range of downstream tasks^5–13^. These models, often implemented as deep language models, adept at data-scarce scenarios where traditional approaches struggle, offering both the improvement on predictive performance and the ability to capture biologically meaningful representations. Specifically, the representations learned during pre-training can be interpreted using post hoc analysis or by-design strategies to uncover valuable biological insights^14^. Fine-tuning expands this capability by enabling these models to adjust their learned representations for specific tasks, generating biological insights that are directly relevant to the given fine-tuning objectives^6,10,15^. By combining broad generalization from pre-training with task-specific adaptability through fine-tuning, pre-trained foundation models are particularly suitable for representing and interpreting the complex dynamics of TRNs, especially in the settings where transcription regulator landscapes are highly incomplete and poorly characterized.

Foundation models pre-trained on DNA sequences have demonstrated remarkable capabilities in deciphering the regulatory architecture of genomes. They effectively capture the organizational syntax of genomic sequences and decode the *cis*-regulatory grammar that governs gene transcription^7,8,11,12,16^. However, these models often fall short in addressing the roles of *trans*-acting transcription regulators, which are essential for a comprehensive understanding of gene transcription. A recent study seeks to bridge this gap by integrating DNA-binding motifs of transcription regulators and chromatin accessibility to improve gene expression predictions^10^, and another study also integrates motifs with multi-omics data to infer gene regulatory architecture^17^. However, these approaches are limited as accessible motifs account for only a fraction of actual binding events, and many transcription regulators lack well-characterized motifs^18,19^. Binding events are often modulated by co-factors, chromatin context, and epigenetic modifications, which extend beyond simple motif recognition, influencing how and where transcription factors bind to their target regions. Consequently, motif-dependent approaches are insufficient to capture the intricate and context-specific interactions among transcription regulators, highlighting the need for foundation models that directly represent and learn these complex interactions.

Here, we introduce ChromBERT, a foundation model specifically designed to directly model genome-wide combinatorial binding patterns of transcription regulators. Pre-trained on the Cistrome-Human-6K dataset, comprising large-scale, heterogeneous ChIP-seq data, ChromBERT effectively captures the interaction syntax of transcription regulators across diverse genomic contexts (Fig. 1a). By leveraging advanced transformer architectures, ChromBERT generates context-specific, biologically interpretable representations of TRNs and their constituent transcription regulators, enabling precise biological interpretation of regulatory roles and the functional collaborations of these regulators. ChromBERT demonstrates superior performance in cistrome imputation for unseen cell types through prompt-enhanced fine-tuning, outperforming previous methods, and showcasing robust generalization across a diverse range of specific cell types. More importantly, when fine-tuned for cell-type-specific downstream tasks, ChromBERT not only achieves superior predictive performance but also adapts its representations to reflect the unique regulatory architecture of each cell type. This adaptability provides clear, interpretable insights into the roles of transcription regulators in driving observed regulatory effects and dynamic changes, without requiring additional cell-type-specific genomic data for each regulator (Fig. 1b). This capability is particularly significant in addressing the challenges posed by sparse transcription regulator data, a common limitation in most biological settings, where comprehensive datasets for transcription regulators are often unavailable.

**Fig. 1.**
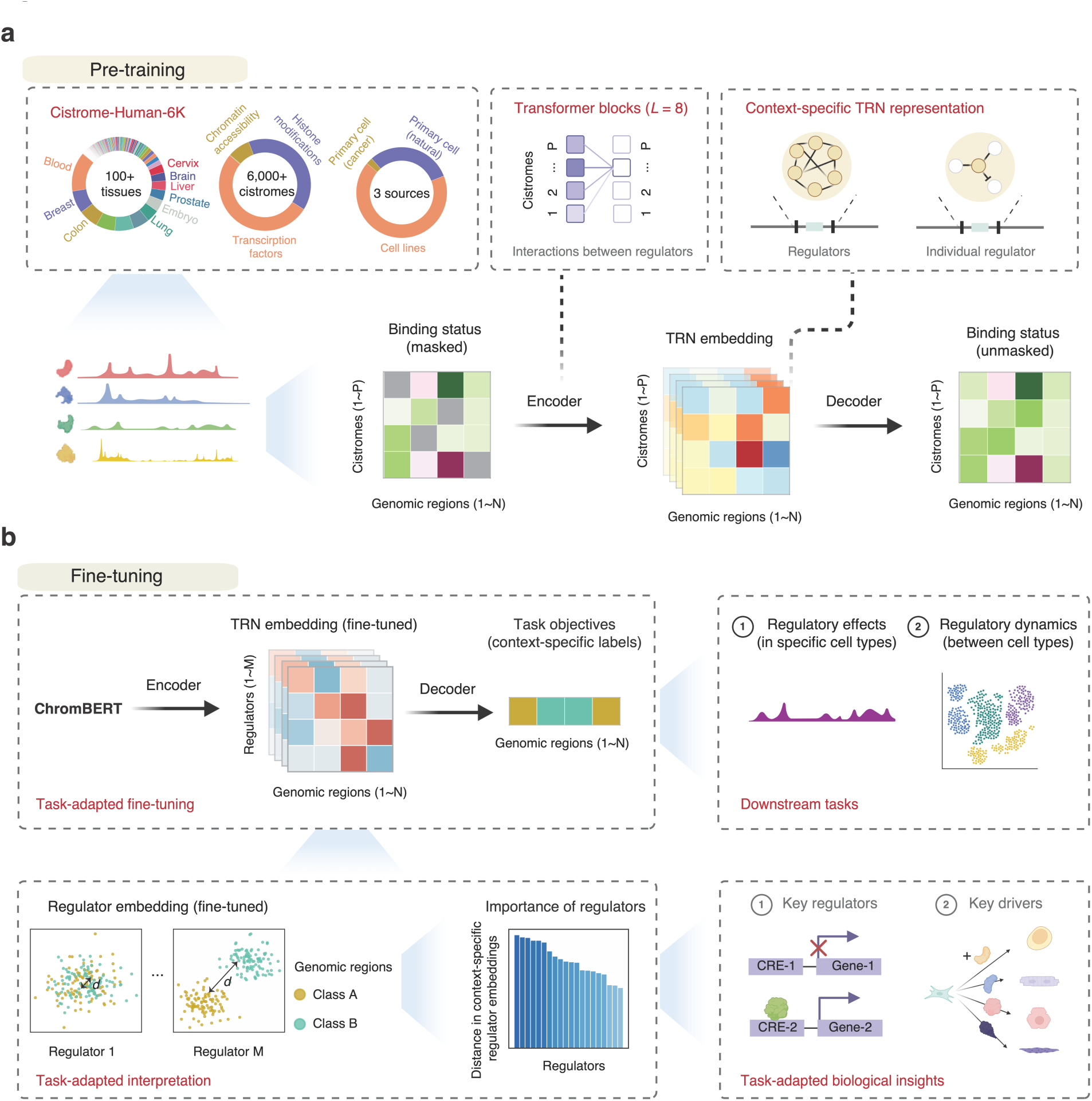
The ChromBERT architecture. **a**. Pre-training schematic of ChromBERT. ChromBERT was pre-trained on the Cistrome-Human-6K dataset, which encompasses cistromes derived from over 100 tissues, sourced from three primary categories: cell lines, primary cancer cells, and naturally occurring primary cells. The input consisted of a binding status matrix, representing the binding statuses of 6,391 cistromes (*Y*-axis in the heatmap, input features) across genome-wide 1-kb intervals (*X*-axis in the heatmap, training samples). The pre-training process employed a masked learning strategy. ChromBERT employed eight self-attention transformer blocks in an encoder to convert masked binding status matrix (where grey in heatmap indicates masked values and other colors denote discretized binding statuses) into TRN representations, complemented by a decoder to reconstruct the binding statuses. During pre-training stage, binding statuses of 15% cistromes were randomly masked, and ChromBERT was trained to predict the masked objectives using the remaining unmasked binding statuses (see Methods for details), which reinforces the model’s ability to capture intricate interaction syntax of these cistromes. ChromBERT generates TRN embeddings (where colors indicate different feature values) representing context-specific TRN, which can be can be further decomposed into regulator embeddings that capture the roles of individual regulators within the TRN. **b.** Fine-tuning schematic of ChromBERT. *Task-adapted fine-tuning*: ChromBERT can be fine-tuned for various transcription regulation-related downstream tasks, such as predicting regulatory effects in specific cell types or regulatory dynamics between cell types. The learning objectives in these tasks are genomic-context-specific labels, which represent the regulatory effects or dynamics of different genomic regions in the given cell type. *Task-adapted interpretation*: Fine-tuning adapts ChromBERT’s TRN embeddings to these task objectives, allowing for the interpretation of transcription regulators’ roles. The key principle is that regulators with significant differences in their embeddings across distinct groups of genomic regions (classified by their regulatory effects) can be identified as key regulators for the task. This approach helps uncover key transcription regulators driving cell-type-specific regulatory effects and mediating cell state transitions.

## Results

### Overview of ChromBERT

ChromBERT distinguishes itself from other genomics foundation models that primarily focus on DNA sequence organization^7,8^ by modeling the combinatorial binding patterns of thousands of transcription regulators and learning their intricate interaction syntax (Fig. 1a). To enable this, ChromBERT was pre-trained on a large-scale corpus of transcription regulator landscapes, leveraging self-attention mechanisms within a transformer architecture^20^ to learn how transcription regulators interact with others at each genomic region. Using a masked learning strategy akin to BERT^21^, the model captures context-dependent interactions of transcription regulators across diverse genomic contexts in a self-supervised manner. These pre-training techniques equip ChromBERT with a foundational understanding of TRNs, enabling strong generalization across a variety of downstream transcription regulation tasks.

The pre-training dataset for ChromBERT was constructed by compiling and filtering nearly all publicly available human ChIP-seq, DNase-seq, and ATAC-seq data from the Cistrome Data Browser^22^ (see Methods for details). This process resulted in the Cistrome-Human-6K dataset, comprising 6,391 qualified cistromes (genome-wide maps of the *cis*-regulatory binding sites of *trans*-acting regulators^23^) for 991 transcription regulators, 76 histone modifications, and chromatin accessibility from various cell types (Fig. 1a, Supplementary Table. S1). Binding signals from all cistromes were discretized into binding statuses, which were then aggregated for each 1-kilobase (kb) genomic interval to construct input features for ChromBERT (Supplementary Fig. S1a, b). This approach enabled the model to learn context-dependent interactions among transcription regulators. Given the limited number of cistromes in individual cell types (Supplementary Fig. S1c), we pooled all qualified human cistromes from different cell types to enrich the diversity of input features. While this cell-type-agnostic strategy introduces variability in representation frequencies for transcription regulators and cell types, it maximizes the use of all available data and supports the model’s ability to generalize. Pre-training on over two million such 1-kb genomic intervals enhanced ChromBERT’s capacity to capture intricate interaction patterns and achieve strong awareness to different genomic contexts. For each genomic region presented to ChromBERT, the pre-trained model embeds it into a TRN embedding that represents the combinatorial interactions among over a thousand transcription regulators within the given region (Fig. 1a). Each regulator’s embedding, encoded as a 768-dimensional vector within the TRN embedding, represents its contextual interactions with other regulators in the same genomic region.

### Pre-trained embeddings reveal human regulatory architecture

After pre-training, we interpreted the pre-trained embeddings from ChromBERT to check whether they could effectively reveal the transcription regulatory architectures specific to their genomic regions. Given the diversity in transcription regulatory architectures across different genomic regions, we hypothesized that pre-trained TRN embeddings would cluster in similar patterns for different groups of regulatory elements. Applying ChromBERT to different groups of regulatory elements on chromosome one, as defined by ChromHMM states^24^, we observed that TRN embeddings generally clustered separately (Supplementary Fig. S2a), suggesting that they effectively delineate distinct regulatory architectures. We also hypothesized that TRN embeddings of nearby genomic regions would display higher similarity due to shared local chromatin structure and transcription regulators. As expected, we noted that the similarity in TRN embeddings decreased with increasing distance between regions (Supplementary Fig. S2b).

Considering the impact of 3D chromatin organization, which segregates the genome into potential functional units^25^, we examined the association between TRN embeddings and 3D genome architecture. Our results showed that regions within the same topologically associating domains (TAD) exhibited higher similarity in their embeddings compared to those separated by TAD boundaries (Supplementary Fig. S2c). Furthermore, long-range genomic region pairs with chromatin contacts showed significantly higher embedding similarity than those without contacts (Supplementary Fig. S2d). To further evaluate how TRN embeddings relate to 3D genome organization, we trained a deep learning model that integrates embedding similarity with Hi-C contact data from long-range genomic region pairs (see Methods for details). This approach successfully imputed high-resolution chromatin contacts (Micro-C) in human embryonic stem cells (hESCs) and demonstrated robust generalization to unseen cell type (Supplementary Fig. S2e). These findings demonstrate that ChromBERT’s pre-trained TRN embeddings effectively represent the complex regulatory architecture of the human genome, providing a powerful framework for characterizing transcriptional regulatory architecture and chromatin organization.

Next, we explored the effectiveness of ChromBERT’s regulator embeddings in representing the functional collaborations among transcription regulators. Each regulator embedding at a given genomic region encapsulates its contextual relationship with other regulators. Intuitively, regulator pairs with strong functional collaborations should exhibit greater embedding similarity. To validate this, we averaged the embeddings of each regulator across genomic regions on chromosome one and calculated the pairwise similarities. This analysis revealed a highly heterogeneous pattern of co-association among transcription regulators, forming distinct clusters (Supplementary Fig. S2f). We identified a specific cluster involving well-characterized transcription factors and co-activators, such as SMARCA4, EP300, BRD4, MED1, FOXM1, and MYC (Supplementary Fig. S2f), indicating functional associations among transcription regulators with high similarity in embeddings.

Moreover, pairs of transcription regulators with high embedding similarity correspond to increased protein-protein interaction frequencies (as defined by affinity-purification mass spectrometry data from BioPlex 3.0^26^) and functional association (as defined by hallmark gene sets and ontology gene sets from the Molecular Signatures Database^27^) (Supplementary Fig. S2g). This provides strong evidence that pre-trained regulator embeddings can reflect the functional collaborations of transcription regulators. Considering the context-specific nature of TRNs, we also examined the embedding similarities of regulator pairs across distinct genomic regions. RNF2, a classical component of the Polycomb Repressive Complex 1 (PRC1)^28^, exhibited high embedding similarity with factors typically associated with transcription repression (such as PCGF2) at some genomic regions, as well as with regulators involved in transcription activation (such as EP300 and MYC) at others (Supplementary Fig. S2h). This pattern largely corresponds to RNF2’s dual functional roles in chromatin^29^. These findings demonstrate the robust ability of ChromBERT to represent context-dependent functional collaborations among transcription regulators.

### ChromBERT boosts cistrome imputation in unseen cell types

Despite extensive efforts in experimental profiling, the genomic landscapes of transcription regulators remain far from comprehensive, particularly given their vast number. To address this limitation, we fine-tuned ChromBERT to impute cistromes of transcription regulators, *i.e.*, to predict the presence or absence of binding events, in previously unseen cell types. A significant challenge in this task lies in effectively integrating cell-type-specific information to facilitate transfer across diverse cell types. Inspired by the advances in prompt-enhanced fine-tuning, a highly efficient and flexible approach that has proven transformative in pre-trained deep language models^30–33^, we adapted this technique for cell-type-specific cistrome imputation. Specifically, we designed cell-type-specific prompts derived from chromatin accessibility profiles or transcriptome profiles to guide the model (see Methods for details). These prompts served as cell-type-specific cues, seamlessly integrating with pre-trained regulator embeddings from ChromBERT to support flexible and accurate predictions for a wide range of transcription regulators across diverse cell types (Fig. 2a). To align these different prompts and embeddings within a unified feature space, we fine-tuned the model using approximately one hundred cistromes from multiple cell types, supplemented by cell-type-specific DNase-seq or transcriptome data (see Method for details). This integration ensured that the model could effectively reconcile variability across cell types and transcription regulators. After this alignment, ChromBERT has the capability to systematically impute cistromes for any given cell type-regulator combination without requiring additional cell-type-specific or regulator-specific fine-tuning.

**Fig. 2.**
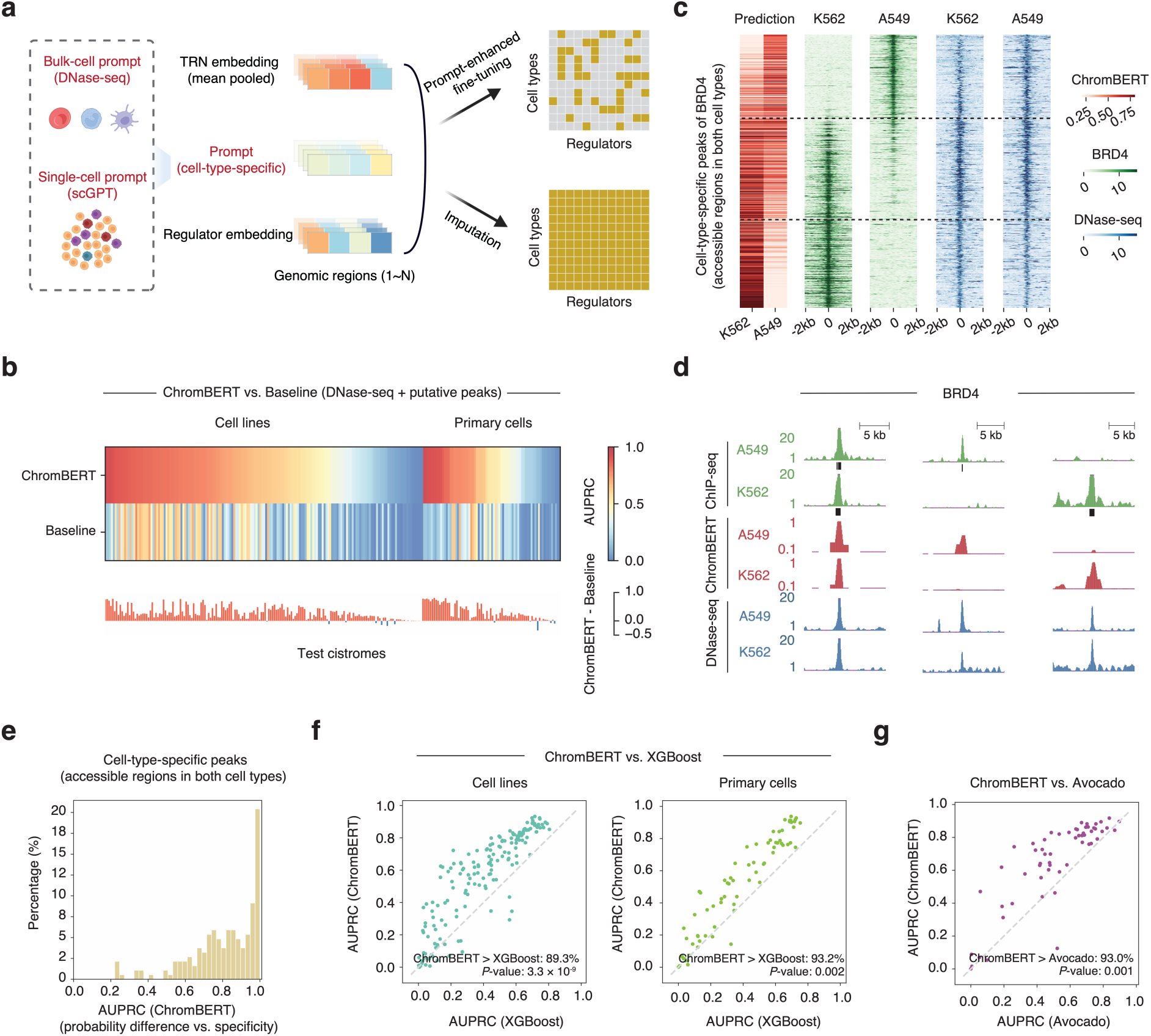
ChromBERT boosts cistrome imputation. **a.** Schematic of cistrome imputation in new cell types or single cells by prompt-enhanced fine-tuning of ChromBERT. This process utilizes TRN embedding along with cell-type-specific prompts and regulator embeddings (see Methods for details). **b.** Performance comparison of cistrome imputation tasks in cell lines and primary cells. The comparison is between ChromBERT with DNase-seq prompts, and a baseline method that utilize DNase-seq signals and regulator-specific putative peaks (see Methods for details). The top heatmap shows performances using the AUPRC metric for both approaches, while the bottom bar plot illustrates performance differences. Red bars indicate ChromBERT’s superiority, while blue bars indicate the baseline method’s advantage. All test cistromes were excluded from the training dataset for prompt-enhanced fine-tuning. **c.** Heatmaps showing predictive probability of ChromBERT with DNase-seq as prompts (red), alongside ChIP-seq signals (green) and DNase-seq signals (blue) on cell-type-specific peaks of BRD4 that are accessible in both K562 and A549 (see Methods for details). The BRD4 ChIP-seq and DNase-seq signals were normalized to the genome background. The black dashed lines divide the test regions into K562 highly-specific, weakly-specific, and A549 highly-specific peaks, arranged from bottom to top. **d.** The UCSC genome browser view illustrates the representative BRD4 binding sites (green) in A549 and K562 cell lines, at consistent peaks (chr17:28,351,650-28,365,238), A549-specific peaks (chr1:220,683,084-220,697,918), and K562-specific peaks (chr3:14,334,778-14,370,222) arranged from left to right. Predictive probabilities of ChromBERT with DNase-seq prompts for BRD4 (red) are shown separately for each cell type, alongside the corresponding DNase-seq signals (blue). **e.** The histogram illustrates ChromBERT’s performance with DNase-seq as prompts on cistrome imputation. Only regions exhibiting cell-type-specific binding events and having similar accessibility in both cell types were evaluated (see Methods for details). The metrics were calculated by using the difference in predictive probabilities between two cell types to predict the ground truth of cell-type-specificity obtained from ChIP-seq peaks. This evaluation included 186 cell-type pairs for 24 regulators. **f.** Scatter plots show the performance comparison between ChromBERT with DNase-seq prompts and XGBoost (see Methods for details) for cistromes in cell lines (left) or primary cells (right). **g.** Scatter plots show the performance comparison between ChromBERT utilizing DNase-seq as prompts and Avocado (see Methods for details). For **f** and **g**, the diagonal gray line indicates equal performance, with percentages of cistromes where ChromBERT outperforms annotated alongside the statistical significance calculated by a two-sided Student’s *t*-test.

ChromBERT demonstrated superior performance in cistrome imputation across a wide range of bulk cell types by leveraging DNase-seq data as prompts. It achieved a mean area under the precision-recall curve (AUPRC) of 0.552 on imputed cistromes, significantly outperforming the baseline approach, which achieved a mean AUPRC of 0.270. The baseline method, which integrates regulator-specific putative peaks and DNase-seq profiles, is a common strategy in the absence of ChIP-seq data. This performance advantage was consistent across both cell lines and primary cells (Fig. 2b). ChromBERT also exhibited exceptional robustness, with its performance largely unaffected by the representation frequency of imputed cell types or transcription regulators in the pre-training dataset (Supplementary Fig. S3a, b). This robustness underscores ChromBERT’s reliability across diverse biological contexts. Notably, one of the most challenging aspects of cistrome imputation is identifying cell-type-specific binding sites, which are biologically significant yet computationally challenging, especially when chromatin accessibility is similar across cell types. ChromBERT demonstrated superior performance in classifying cell-type-specific binding sites that could not be explained by differential chromatin accessibility alone (Fig. 2c-e). To further validate its effectiveness, ChromBERT was compared to an XGBoost model trained directly on the same input cistromes used during pre-training. ChromBERT consistently outperformed XGBoost (Fig. 2f), underscoring the predictive power of its pre-training approach. Furthermore, when benchmarked against Avocado, a state-of-the-art model for cistrome imputation^34^, ChromBERT achieved superior accuracies across applicable test datasets (Fig. 2g).

Imputing single-cell cistromes is significantly more challenging than bulk-cell cistromes due to the limited availability of paired single-cell omics data^35,36^. Despite the rapid expansion of single-cell transcriptomics over the past decade, the lack of corresponding transcription regulator data impedes our understanding of transcription regulation at the single-cell level, where cell-type and state-specific dynamics play crucial roles. Building on the successful alignment of bulk-cell RNA-seq-based prompts from scGPT^6^ with pre-trained ChromBERT embeddings for bulk-cell cistromes imputation (Supplementary Fig. S4a), we evaluated its potential to impute cistromes at single-cell resolution by replacing bulk-cell RNA-seq prompts with single-cell prompts (see Method for details). We evaluated ChromBERT’s single-cell imputation performance using a human peripheral blood mononuclear cells (PBMCs) single-cell multi-omics dataset. Two key aspects were assessed: predictive accuracy in B cells, using ChIP-seq data from GM12878 as ground truth, and the single-cell-specificity of prediction, using motif accessibility inferred from single-cell ATAC-seq data as ground truth (see Methods for details). ChromBERT achieved significantly higher predictive performance for eight regulators in B cells, outperforming a combination of DNA-binding motifs and imputed chromatin accessibility (Supplementary Fig. S4b). Additionally, ChromBERT exhibited robust single-cell-specific performance (Supplementary Fig. S4c, d). These results underscore the utility of ChromBERT in cistrome imputation and highlight its potential to extend transcription regulatory studies by leveraging diverse prompts tailored to a range of biological settings.

### ChromBERT reveals key regulators in specific cell types

Transcription regulation is highly complex, as regulatory effects vary across both cell types and genomic contexts. Within each cell type, genomic regions exhibit specific regulatory effects (e.g., active enhancers or promoters), but the roles of transcription regulators mediating these behaviors are often unclear due to limited cell-type-specific data. To address this, we fine-tuned ChromBERT for cell-type-specific tasks, enabling it to learn regulatory effects (such as enhancer activity or gene expression level) for diverse genomic regions within each cell type (Fig. 3a). By interpreting these fine-tuned TRN representations, ChromBERT offers insights into the roles of transcription regulators in mediating cell-type-specific regulatory effects. Specifically, a regulator’s contextualized embedding can be interpreted to reflect its context-dependent roles across genomic regions^15,37^. When the differences between a regulator’s embeddings for distinct groups of genomic regions correspond to observed regulatory effects, the regulator can be identified as a key contributor in the biological context of interest (see Method for details). This approach is particularly beneficial for rare or hard-to-study samples with limited experimental profiling. We fine-tuned ChromBERT for three downstream tasks to demonstrate its ability.

**Fig. 3.**
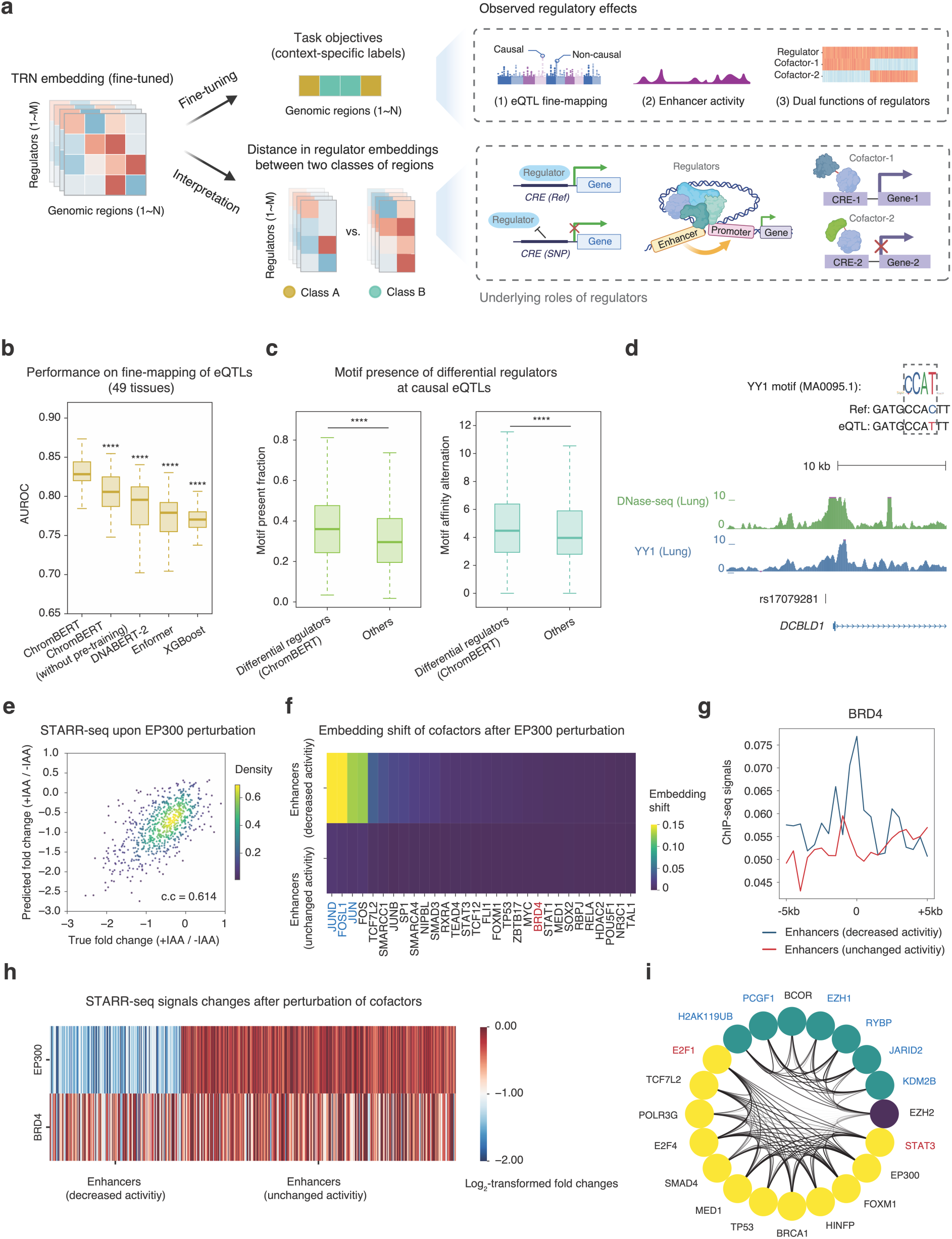
ChromBERT reveals key regulators in cell-type-specific regulatory effects. **a**. Schematic of ChromBERT’s fine-tuning in modeling cell-type-specific regulatory effects and interpreting the underlying roles of transcription regulators. From left to upper right, ChromBERT is fine-tuned to generate fine-tuned TRN embeddings (colors in the heatmap indicate feature values) to predict observed regulatory effects in specific cell types or tissues, where the task objective is context-specific labels across diverse genomic loci. From left to lower right, fine-tuned TRN embeddings are interpreted to reveal the roles of transcription regulators corresponding to observed regulatory effects (see Methods for details). **b.** Box plots comparing the performance in classifying causal versus non-causal eQTLs across 49 human tissues for ChromBERT, ChromBERT (without pre-training), DNABERT-2, Enformer, and XGBoost, as indicated by AUROC (see Methods for details). **c.** Box plots comparing the motif presence fraction and motif affinity alternation between differential regulators (top 20% showing the highest embedding shifts between two groups of eQTLs in each tissue) and other regulators. The motif presence and affinity (log-odds scores) around +/-10bp causal eQTLs were determined using FIMO (v5.5.5)^83^. For each regulator, motif presence was identified by p-value < 0.01 in either wild type or mutant form, and only sites with motif affinity > 0 in either wild type or mutant form were analyzed for motif affinity alternation. For **b** and **c**, the statistical significance was performed by a two-sided Mann-Whitney *U*-test and **** represents *p*-value < 1 × 10^−4^. The center lines mark the median, the box limits indicate the 25th and 75th percentiles, and the whiskers extend to 1.5× the interquartile range from the 25th and 75th percentiles. **d.** The UCSC genome browser view shows YY1 signals (blue) near eQTL rs1707928, along with DNase-seq signals in lung (green). The YY1 ChIP-seq signals were from GSE32465^84^, and DNase-seq signals were from GSE18927^79^. The DNA sequence in reference genome and genetic variants around the eQTL were shown in the upper right, and the YY1 binding motif from JASPAR was also shown. **e.** Scatter plots showing the ground truth of log_2_-transformed fold change of STARR-seq signals upon EP300 perturbation and predictions by ChromBERT at test enhancers. The Pearson’s correlation coefficient is annotated in the plot, and the color represents the density of points. STARR-seq signals were from previous study (GSE156741^44^). **f.** Heatmap showing the distance in regulator embeddings of potential cofactors of EP300 at the test enhancers before and after in silico perturbation for EP300/CREBBP. Two groups of enhancers were depicted: those with decreased activity (log_2_-transformed fold change < −1, *n* = 164) and those unchanged (−0.5 < log_2_-transformed fold change < 0.5, *n* = 343) after IAA-treated EP300/CREBBP depletion. Potential cofactors of EP300 were defined as top 50 regulators that showing the highest embedding similarity with EP300 at all test enhancers. The color represents the shifts (1 - cosine similarity) in embeddings. Representative regulators showing high embedding shift at decreased enhancers are highlighted in blue and representative regulators showing low embedding shift at decreased enhancers are highlighted in red. **g.** Line charts showing average ChIP-seq profile of BRD4 around test enhancers. BRD4 ChIP-seq data was from previous study (GSE57628^80^). **h.** Heatmap showing log_2_-transformed fold changes of STARR-seq signals after IAA-treated EP300/CREBBP or BRD4 depletion, test enhancers were divided into two groups with decreased or unchanged activity after IAA-treated EP300/CREBBP depletion. **i.** Circos plot showing embedding similarities with EZH2 at classical (w/ H3K27me3) and non-classical (w/o H3K27me3) genomic loci, highlighting two groups of regulators having locus-preferential embedding similarities with EZH2. The classical group shows higher embedding similarity with EZH2 at classical loci compared to non-classical loci (embedding similarity with EZH2 at classical loci ranking the top 5% among all regulators and the embedding similarity difference between classical loci and non-classical loci > 0.1), and the non-classical group exhibits the converse pattern. The classical group was marked in green and the non-classical group was marked in yellow. The known cofactors in classical function of EZH2 were highlighted in blue, and the known cofactors in non-classical function of EZH2, E2F1 and STAT3, were highlighted in red. Each node represents a regulator, and the transparency of edges linking two nodes represent the embedding similarity of two regulators, only edges with high pairwise embedding similarity (> 0.8) were plotted.

The first task involved tissue-specific fine-mapping of expression quantitative trait loci (eQTL), a specific type of genetic variants that influence the expression levels of genes. Fine-mapping within eQTLs helps to pinpoint the exact causal variants affecting gene expression, which is crucial for understanding the genetic basis of complex traits and diseases^38^. To address this, we fine-tuned ChromBERT to classify causal and non-causal variants using the latest data from the eQTL Catalogue. We incorporated DNA sequence variation by integrating a DNA sequence prompt from DNABERT-2 into ChromBERT (see Methods for details). As shown in Fig. 3b, fine-tuned ChromBERT achieved a mean area under receiver operating characteristic (AUROC) of 0.828, outperforming Enformer (mean AUROC = 0.770), DNABERT-2 model alone (mean AUROC = 0.788), ChromBERT without pre-training (mean AUROC = 0.804), and an XGBoost model across 49 human tissues. Fine-tuned ChromBERT generated contextualized embedding for transcription regulators across eQTLs, allowing the identification of differential regulators that exhibited the significant embedding distance between casual and non-causal eQTLs (see Methods for details).

Chromatin accessibility, indicated by DNase-seq signals, varied the most on average across 49 tissues (Supplementary Fig. S5a), aligning with its known significant role in transcription regulation^39,40^. Further analysis of public DNase-seq profiles across 10 tissues confirmed that causal eQTLs exhibited elevated chromatin accessibility (Supplementary Fig. S5b). Additionally, we also investigated transcription factors that exhibited significant distance in contextualized embeddings between the two groups of eQTLs (Supplementary Fig. S5b). These differential factors not only have a higher motif presence at causal QTLs but also demonstrated significant alternations in motif affinity following single-nucleotide genetic variations at these loci (Fig. 3c). For instance, in a case study of causal eQTL rs17079281 influencing the transcription of *DCBLD1* in lung, we found that a C to T variation led to increased motif affinity of YY1 (Fig. 3d), a differential regulator identified by ChromBERT. It is consistent with a previous study demonstrating that rs17079281 decreases lung cancer risk by creating an YY1-binding site to suppress *DCBLD1* expression^41^. These results indicate that ChromBERT is effective in fine-mapping eQTLs, providing valuable insights into the genetic variants that drive gene expression changes.

The second task was to fine-tune ChromBERT for modeling genome-wide enhancer activity quantified from STARR-seq^42^. In this task, ChromBERT showed superior prediction performance than Enformer, DNABERT-2, ChromBERT without pre-training, and XGBoost model in predicting STARR-seq signals in HCT116 cells (Supplementary Fig. S5c). However, although numerous transcription regulators and cofactors are known to co-bound at active enhancers, perturbation experiments for regulators suggests that differential cofactor dependencies define distinct types of human enhancer^43,44^. To explore the cofactor dependencies in enhancer activity, we further fine-tuned ChromBERT to model perturbation effects of regulators using a STARR-seq dataset for EP300/CREBBP, BRD2 and CDK7 perturbations^44^. The model performed well in predicting both wild-type and perturbed STARR-seq signals, capturing at least a fraction of perturbation effects (Fig. 3e, Supplementary Fig. S5d-f). The perturbation model enables us to capture key regulators dependent on perturbated regulators. We interpreted regulators with significant embedding shifts upon EP300 perturbation at enhancers with decreased activity as key cofactors of EP300 (see Methods for details). This approach identified cofactors such as FOSL1 and JUND having strong functional associations with EP300 (Fig. 3f), which are consistent with the findings of the previous study^44^. Interestingly, while other well-characterized cofactors like BRD4 were also enriched at these regions, they did not show a close association with EP300 in our interpretation analysis (Fig. 3f, g). To validate this finding, we re-analyzed STARR-seq data following BRD4 perturbation and confirmed the weak functional association between BRD4 and EP300 in enhancer activities (Fig. 3h). These results demonstrate that fine-tuning ChromBERT can help to investigate the complex regulatory mechanisms governing enhancer activity.

The third task was to distinguish distinct functional loci of a multi-function transcription regulator. EZH2, a core component in Polycomb Repressive Complex 2 (PRC2)^45^, has been reported to have two context-dependent functions: a classical function involving H3K27me3 and a non-classical function independent of H3K27me3^29,46^. We fine-tuned ChromBERT to classify classical and non-classical sites of EZH2, where ChromBERT showed a better performance in this task than other methods (Supplementary Fig. S5g). Further interpretation of the fine-tuned representations revealed their functional collaborations with EZH2 in determining EZH2’s context-dependent functions (see Methods for details). Two distinct groups of transcription regulators showed high co-association with EZH2 (Fig. 3i). The first group is strongly associated with regulators involved in repressive function, aligning with the classical function of EZH2. In contrast, the second group is related to active function, including E2F1 and STAT3, two known cofactors in EZH2’s non-classical function^29,47,48^. These results demonstrate the ability of ChromBERT to decipher the context-dependent functions of transcription regulators.

### ChromBERT uncovers key drivers in cell state transition

Understanding the transcription regulatory dynamics underlying cellular heterogeneity and cellular reprogramming is crucial for gaining deep insights into development, disease processes, and drug discovery. To investigate these dynamics and identify key regulators involved in cell state transitions, we fine-tuned ChromBERT to model genome-wide dynamics of regulatory effects, *i.e.*, to capture transcriptomic and chromatin accessibility changes, between two cell types. By interpreting the fine-tuned representation adapted to these tasks, we identified driver regulators responsible for the transition between two cell states (Fig. 4a; see Methods for details).

**Fig. 4.**
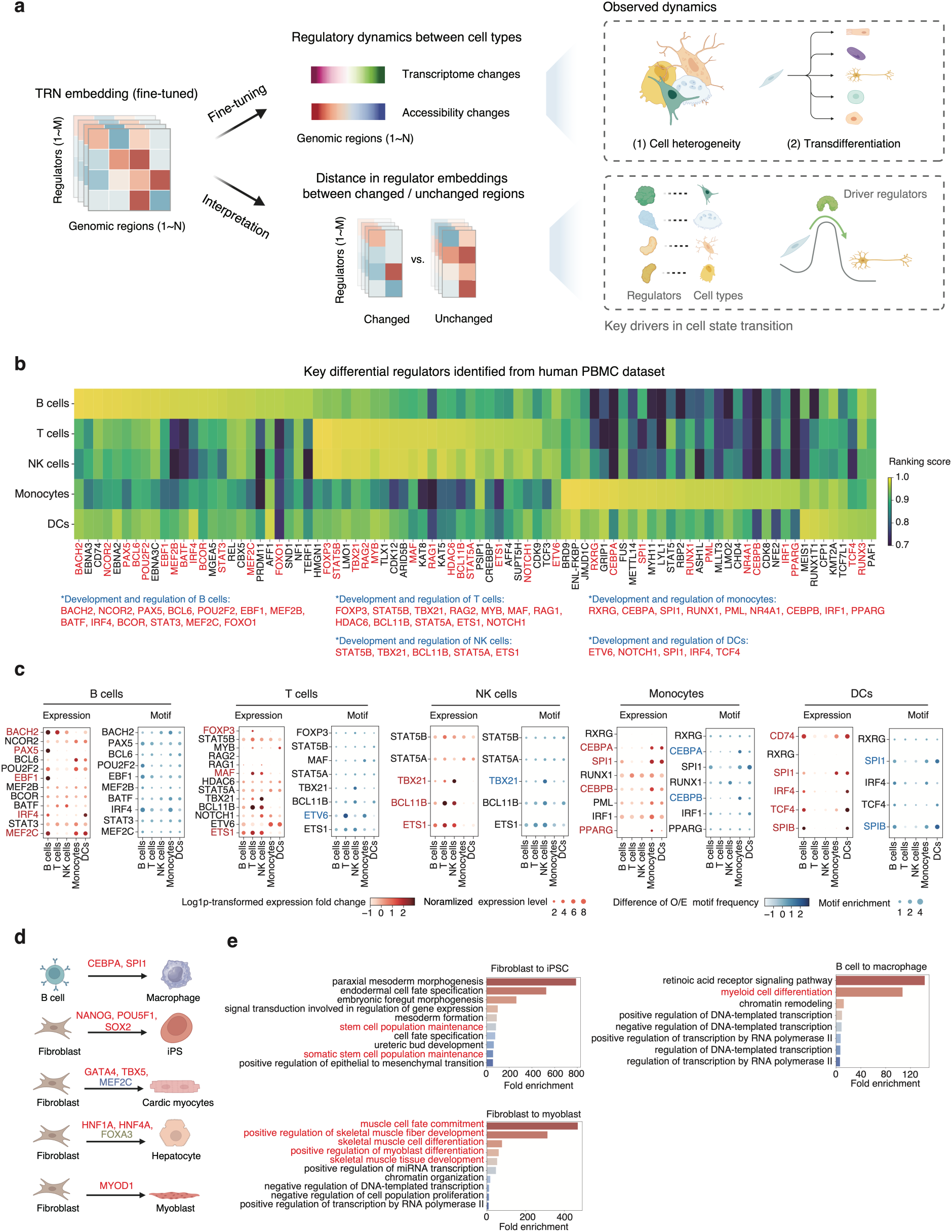
ChromBERT uncovers key regulators in cell state transition. **a**. Schematic of the fine-tuning of ChromBERT in modeling regulatory dynamics between cell types and interpreting underlying roles of transcription regulators. From left to upper right, ChromBERT is fine-tuned to generated fine-tuned TRN embeddings (colors in heatmap indicate feature values) to predict observed regulatory dynamics between cell types, where the task objective is genome-wide changes in transcriptome or chromatin accessibility. From left to lower right, fine-tuned TRN embeddings are interpretated to reveal roles of transcription regulators corresponding to observed regulatory dynamics during cell state transition (see Methods for details). **b**. Heatmap showing cell-type-specific ranking scores of all key regulators identified in five major cell types: B cells, T cells, NK cells, monocytes and DCs. The cell-type-specific ranking scores evaluate the importance of regulators in differentiating the given cell type from all other cell types (see Methods for details). All key regulators identified were generally classified into four categories, B cells-specific, T cells/NK cells-specific, monocytes-specific and DCs-specific according to the cell-type-specific ranking scores for all 84 identified key regulators. Reported key regulators in each cell type were highlighted in red and specifically annotated below the heatmap. **c**. Dot plots showing the low coverage of reported important regulators in each cell type by cell-type-specific differential expression analysis or motif analysis. Reported key regulators covered by differential expression analysis were highlighted in red, while those covered by differential motif analysis were highlighted in blue. Regulators showing differential expression are those with notably higher expression in a specific cell type compared to others (log1p-transformed expression level in the given cell type − average expression level in other cell types > 1). Regulators showing differential motif enrichment are those with notably higher motif enrichment at regions with increased chromatin accessibility in the specific cell type compared to others (observed/expected motif occurrence frequency in the given cell type – average observed/expected motif occurrence frequency in other cell types > 1). The sizes of points indicate the expression level or motif enrichment, and the colors indicate the difference between the given cell type and mean value in other cell types. **d**. The extent to which ChromBERT recapitulates known driver regulators across five transdifferentiation processes. Known driver regulators recapitulated by ChromBERT were highlighted in red. MEF2C, not recapitulated by ChromBERT, was marked in blue. And FOXA3, lacking related cistromes in our model, was marked in brown. **e**. Gene ontology (GO) enrichment analysis for 25 identified key regulators using DAVID online tool (https://david.ncifcrf.gov/tools.jsp)^85^. The top ten human gene ontology biological process (BP) terms with a false discovery rate (FDR) < 0.05 and high fold enrichment were displayed. The GO terms involved in the terminal cell states were highlighted in red.

We first examined the transcription regulatory dynamics within the human PBMC dataset, which includes five cell types: B cells, T cells, natural killer (NK) cells, dendritic cells, and monocytes, based on marker genes. ChromBERT was fine-tuned to predict the genome-wide changes in the transcriptome or chromatin accessibility between these cell types in a pairwise manner (see Methods for details), enabling the model to capture the dynamics of TRNs. In this regression task, ChromBERT outperformed other methods (Supplementary Fig. S6). To evaluate the impact of transcription regulators, we analyzed the embedding distance between two groups of genomic regions characterized by increased or unchanged gene transcription or chromatin accessibility between two cell types. This interpretation analysis revealed the contribution of specific regulators to cellular heterogeneity and identified key regulators unique to each cell type (see Methods for details). Notably, 37 out of 84 identified key regulators have previously been reported to play important functions in the corresponding cell types (Fig. 4b). For instance, BACH2, NCOR2, PAX5, BCL6, BCOR and IRF4 are well-characterized factors involved in the transcription regulation of memory B cell differentiation^49^. Similarly, FOXP3 and TBX21 are known regulators in T cell lineage commitment^50,51^, CEBPA, CEBPB, and SPI1 are well-characterized regulators in monocytes^52–54^. Strikingly, many of these known functional regulators recapitulated by ChromBERT could not be detected through differential expression analysis or motif enrichment analysis in differentially accessible chromatin (Fig. 4c), two commonly used methods for key regulator identification^55^. Additionally, SCENIC+, a computational framework for single-cell multi-omics inference of gene regulatory networks^56^, also failed to predict most known functional regulators in this dataset (Supplementary Fig. S7a). Remarkably, these known functional regulators displayed greater distances and higher rankings in embeddings in the fine-tuned model than the pre-trained model (Supplementary Fig. S7b, c), confirming the effectiveness of fine-tuning and representation adaption in uncovering important regulators in cellular heterogeneity. These findings suggest that ChromBERT can accurately capture the dynamics of TRNs between different cell types and highlight important transcription regulators.

Next, we focused on transdifferentiation, which involves directly reprogramming from starting cells to target cells. In this scenario, ChromBERT was fine-tuned to predict genome-wide transcriptome and chromatin accessibility changes across five transdifferentiation processes. ChromBERT consistently outperformed other methods (Supplementary Fig. S8a, b). Driver regulators were identified by ranking the embedding distances between genomic regions with increased or unchanged transcription or chromatin accessibility during transitions from starting to target cell states (see Methods for details). Most known driver regulators involved in these transitions were successfully recapitulated by this interpretation analysis (Fig. 4d, Supplementary Fig. S8c). Moreover, the remaining identified key regulators showed higher embedding similarity with known driver regulators (Supplementary Fig. S8d), indicating that newly identified transcription regulators may function collaboratively with known drivers. For instance, identified key regulators in three cell state transition processes, from fibroblast to iPSCs, from B cells to macrophage, and from fibroblast to myoblast, were highly enriched in gene ontology (GO) terms related to the target cell fate determination (Fig. 4e). Additionally, we observed that these known driver regulators have higher rankings and more significant shifts in regulator embeddings in the fine-tuned model than the pre-trained model (Supplementary Fig. S8e, f), also confirming the effectiveness of fine-tuning and representation adaption. These results demonstrate ChromBERT’s ability to identify key transcription regulators driving transdifferentiation processes, providing valuable insights for targeted cell fate manipulation.

## Discussion

An essential aspect of studying transcription regulation is elucidating context-specific TRNs across diverse genomic contexts and cell types. These TRNs are crucial for explaining differences in gene expression, cellular functions, and phenotypes. This study introduces ChromBERT, a deep language model pre-trained on a large-scale human ChIP-seq dataset to directly model genome-wide combinatorial binding patterns of transcription regulators. Through fine-tuning, ChromBERT demonstrates superior performance in a variety of downstream tasks related to transcription regulation, including cistrome imputation in previously unseen cell types or single cells, as well as modeling cell-type-specific regulatory effects and dynamics. In addition, by interpreting transformers-encoded TRN embeddings, ChromBERT sheds light on the roles and functional collaborations of transcription regulators in regulating gene expression and chromatin states. Notably, ChromBERT adapts these embeddings to the specific biological contexts associated with downstream objectives, offering cell-type-specific insights into the regulatory architectures without requiring cell-type-specific genomic data on transcription regulators.

ChromBERT’s architecture and training paradigm offer significant opportunities for expanding its applications to additional downstream tasks in transcription regulation. For example, ChromBERT could play a transformative role in single-cell omics studies by identifying context-specific transcriptional regulatory architectures and their dynamics across heterogeneous cell populations. It could also enhance efforts to understand tissue-specific regulatory mechanisms, aiding studies that investigate the interplay between transcription factors, chromatin states, and lineage-specific differentiation processes. Although ChromBERT primarily focuses on the interactions of transcription regulators within each genomic context, integrating its TRN embeddings from multiple genomic contexts with long-range genomic distances could address emerging challenges, such as enhancer-promoter interaction prediction and the dissection of long-range chromatin interactions. ChromBERT’s potential can be further amplified through integration with other foundation models via prompt-enhanced fine-tuning. By leveraging prompts that encode biological information, ChromBERT can synergistically interact with DNA-sequence-based foundation models, which focus on DNA sequence grammar, or single-cell foundation models, which specialize in single-cell transcriptomics. For instance, prompts can guide the integration of DNA sequence features from DNA-sequence-based foundation models with ChromBERT’s TRN embeddings, enabling joint analyses of sequence and regulatory patterns. Similarly, combining ChromBERT with single-cell foundation models through prompts could yield insights into the gene-regulatory networks underlying single-cell transcriptomic data, enhancing the understanding of dynamic cell states and transitions.

Despite its promising potential, ChromBERT has limitations that hinder its full utility in modeling transcriptional regulation. These challenges arise primarily from the diversity and representation frequency of transcription regulators during pre-training and the complexities of aligning different foundation models for downstream tasks. ChromBERT relies on publicly available cistromes for approximately one thousand transcription regulators, but many regulators lack such data, limiting its representation of the full transcriptional regulatory spectrum. Additionally, the scarcity of cell-type-specific cistromes necessitates pooling data across cell types, which, while broadening regulator inclusion, compromises cell type specificity. Expanding ChIP-seq datasets and incorporating cell type information as metadata could enable ChromBERT to analyze transcription regulation in a cell-type-specific manner and leverage cell-type-specific prompts for downstream tasks. Integration with other foundation models poses further challenges due to differences in latent spaces across modalities, such as DNA sequences and single-cell transcriptomics. For example, accurately predicting variant effects requires better alignment between ChromBERT embeddings and sequence-based embeddings, as the current model struggles to distinguish distinct single-nucleotide variants within the same genomic region. A more comprehensive solution involves pre-training a multi-modal foundation model from scratch, combining diverse data types like DNA sequences, long-range chromatin interactions, and gene expression profiles. This approach could provide a unified framework for transcription regulation, offering systematic and comprehensive insights while overcoming the limitations of post hoc model alignment.

## Methods

### Assembly and binding status encoding of cistromes

#### Assembly of cistromes in Cistrome-Human-6K

To create a comprehensive pre-training corpus, we gathered data from a wide range of cell types using publicly available sources. Initially, we collected all human ChIP-seq, DNase-seq, and ATAC-seq data (cistromes) from the Cistrome Data Browser^22^. We then applied several filtration steps to select qualified datasets:

- Quality control metrics: We included datasets that met at least four out of five quality control metrics provided by the Cistrome Data Browser: sequence median quality score ≥ 25, uniquely mapped ratio ≥ 60%, PCR bottleneck coefficient ≥ 80%, fraction of reads in peaks ≥ 1% and number of 10-fold confident peaks ≥ 500.
- Peak counts: We recorded datasets with more than 100 called peaks for transcription factors and over 1000 called peaks for other transcription regulators.
- Genome coverage: We ensured that more than 60% of genome-wide regions without called peaks were covered by reads.
- Replicate removal: To ensure accuracy in our annotations, we followed a structured approach to associate datasets with cell populations. Specifically, when a dataset explicitly included a cell line annotation, we used the cell line as the primary label. For datasets without cell line annotations, we assigned labels based on a combination of cell type and tissue to avoid conflating datasets with similar cell types derived from different tissues. This approach ensures consistent and biologically meaningful categorization. To avoid redundancy, we retained the highest-ranking dataset for each transcription regulator in the same cell line / cell type, based on the combined score of all quality control metrics. Given the inherent heterogeneity of tissues, all datasets derived from tissues that met our quality control criteria were retained for analysis.

Following these filtration steps, we assembled a dataset Cistrome-Human-6K consisting of 6,391 ChIP-seq, DNase-seq and ATAC-seq cistromes. Additionally, we manually curated its metadata to ensure its accuracy (Supplementary Table. S1). The dataset was subsequently processed into a signal matrix, *S* ∈ ℝ^*N*×*P*^, where each element *S*_*i,j*_ represents average signal (reads per million mapped reads, RPM) of cistromes *c*_j_ ∈ {*c*_1_, *c*_2_, ⋯, *c*_*P*_} (*P* = 6,391) at the *i*-th bin *b*_*i*_ ∈ {*b*_1_, *b*_2_, ⋯, *b*_*N*_} in the genome. The set {*b*_1_, *b*_2_, ⋯, *b*_*N*_} denotes consecutive 1-kb bins split from the human build genome hg38. To avoid potential sequencing bias, blacklist regions from ENCODE were excluded from the analysis (https://www.encodeproject.org/files/ENCFF356LFX/). To focus on co-binding events at genomics regions, only bins with peak presence of more than two cistromes among all cistromes were kept in the analysis. Ultimately, a total of 2,137,894 1-kb bins were retained in the matrix.

#### Binding status encoding of cistromes

Transcription regulators often co-bind to chromatin and function in a combinatorial manner^1^. In ChromBERT, we model the combinatorial binding patterns of cistromes to generate representations for context-specific TRNs. Therefore, the input into ChromBERT is the binding statuses of all assembled cistromes at each individual 1-kb bin *b*_*i*_, which were processed from signal vector *S*_*i*_ = [*S*_*i*,1_, *S*_*i*,2_, ⋯, *S*_*i*,*P*_] at genomic bin *b*_*i*_.

To ensure the comparability and suitability of signals from different cistromes for input into the deep language model, we first discretized these signals into distinct categories representing different binding statuses. Specifically, for each cistrome *c*_j_, we categorized its signals across the entire genome into two groups, “binding” or “non-binding”. The threshold was defined as the 10-th percentile of signals from 1-kb bins with peak presence. Peak files were downloaded from Cistrome Data Browser^22^. This categorization is designed to highlight the biologically meaningful but sparse binding events of transcription regulators in the genome. The “binding” category was further divided into three equal portions, “slightly positive”, “moderately positive” and “strongly positive”, based on the signals within this group. Similarly, the “non-binding” group was divided into two equal portions, “slightly negative” and “strongly negative” (Supplementary Fig. S1a). This finer classification aims to incorporate the impact of binding strength in modeling the co-association pattern of transcription regulators. After binding status categorization for all cistromes, the signal vector can be converted to a binding status vector *X*_*i*_ = [*X*_*i*,1_, *X*_*i*,2_, ⋯, *X*_*i*,*P*_], where *X*_*i,j*_ corresponding to the defined five categories of binding statuses from “strongly negative”, “slightly negative”, “slightly positive”, “moderately positive” to “strongly positive”.

### Architecture and pre-training of ChromBERT

#### The architecture of ChromBERT

ChromBERT architecture includes three parts: (1) an input embedding layer to generate input embeddings from input features; (2) eight transformer blocks (encoder); and (3) output heads adapted to different tasks (decoder).

The input embedding layer converts the binding status vector of each 1-kb bin into a high-dimension representation. The binding status vector in each 1-kb bin, *X*_*i*_, is analogous to a sentence in natural language processing, where binding status of each cistrome, *X*_*i,j*_, is considered as a word in the sentence. Therefore, we utilize the similar input representation approach in BERT model^21^, to generate input embeddings from *X*_*i*_. The input embeddings include position embeddings that represent different cistromes and token embeddings that represent different binding statuses. Each word in *X*_*i*_ is assigned a unique integer position identifier based on its originating cistrome, resulting in a position vector for all words, denoted as 𝑣 = [*id*(1), *id*(2) ⋯, *id*(*P*)]. And token vector is *X*_*i*_, where each item *X*_*i,j*_ belongs to one of five binding status categories. Then ChromBERT employ the commonly used embedding layers in PyTorch (v2.0.1)^57^, *Emb_position_* and *EMb*_*token*_, for two vectors respectively, to map each position and token to a fixed-length embedding vector of dimension *d*_*input*_ = 768. Consequently, the resultant summarized input embeddings for each bin *b*_*i*_, 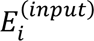 ∈ ℝ^*P*×*dinput*^, is defined as:

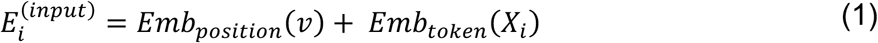

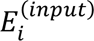 represents the binding statuses of *P* cistromes at the same genomic bin, facilitating the following modeling of their co-association patterns with transformers.

ChromBERT employs a transformer encoder to convert input embeddings 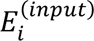 into TRN embeddings 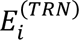 ∈ ℝ^*P*×*d_model_*^, which represents context-specific TRN, where *d_model_* = 768. The encoder comprises eight self-attention transformer blocks^20,21^, each consisting of a multi-head self-attention layer and a feed forward neural network layer (Supplementary Fig. S1b). These self-attention mechanisms operate on the input embeddings, allowing the model to capture intricate interactions among all cistromes. The self-attention mechanism of *l*-th block operates as follows:

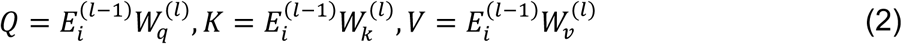

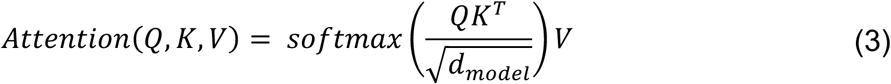

Here, 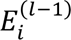 is the output of the (*l* − 1)-th block, and 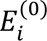 is 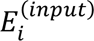, *Q*, *K* and *V* are the query, key and value vectors respectively, each with dimension *P* × *d_model_*. The weight matrices 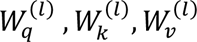 are of dimensions *d_model_* × *d_model_*, and 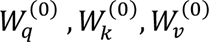 are of dimensions *d_input_* × *d_model_*. Additionally, ChromBERT splits the computation of self-attention into eight attention heads to allow the model to jointly attend to information from different representation subspaces at different positions. To enhance processing efficiency for the large sequence length (*P* = 6,391), ChromBERT incorporates FlashAttention-2 (v2.3.2)^58^, which optimizes attention computation through improved parallelism and work partitioning.

Due to the lack of explicit ground truth for genome-wide combinatorial interactions among transcription regulators, ChromBERT’s pre-training is performed in a self-supervised manner. This involves a decoder architecture that follows the encoder to reconstruct input binding status vector *X*_*i*_ from the TRN embedding 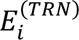, producing an output *Ŷ_i_* ∈ ℝ^*P*×5^. The model setup and training processes are facilitated using PyTorch, ensuring robust configuration, data handling, and computational efficiency.

#### Pre-training with masked learning

ChromBERT employs a masked learning approach, widely used in pre-trained models, to enhance the generalizability of fundamental knowledge acquired during pre-training, which allows for efficient and accurate transfer learning in a variety of downstream fine-tuning tasks^5,6,8,21,59^. In the pre-training of ChromBERT, 15% of the binding statuses in the input sequence *X*_*i*_ were randomly selected. The model’s pre-training objective is to predict these masked statuses based on the context provided by the remaining cistromes. For these 15% binding statuses, we applied a similar masked learning strategy used in BERT^21^. In details, we replaced the binding statuses *X*_*i,j*_ with (1) the [*MASK*] in 80% of the time or (2) a random binding status in 10% of the time or (3) the unchanged *X*_*i,j*_ in 10% of the time. Through this process, ChromBERT gained deep insights into the grammar that governs the combinatorial interactions among transcription regulators, effectively learning to infer masked elements.

Due to the severe class imbalance among the five binding status categories, focal loss^60^ was employed to refine the training process. This loss function modifies the standard cross-entropy loss to focus more on difficult-to-classify instances by decreasing the weight of easily classified examples. In the pre-training stage, only masked objectives were taken account in the loss computation. Let *y*_j_ = *Y*_*i,j*_ denote as the true binding status category of the cistrome *c*_j_ at the *i*-th bin *b*_*i*_, then the focal loss function is defined as:

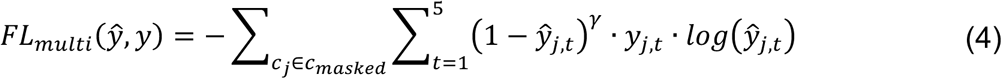

In this formula, *γ* is a focusing parameter, set to 2 by default, which scales the contribution of each example based on the ease of its classification. *y_j,t_* is an indicator variable equals to 1 if *X_*i,j*_ = *t*, and equals to 0 for otherwise, where *t* is one of the five binding status categories. *ŷ*_j,t_* is the predicted probability for the binding status cistrome *c_j_* being *t*. This adjustment ensures that the model pays greater attention to learning from challenging cases, thereby addressing the imbalance in the training data effectively.

Specific hyperparameters during the pre-training of ChromBERT were empirically selected as follows. The learning rate scheduler was configured to a linear schedule, incorporating a warm-up period that comprised 10% of all learning steps, increasing from 0 to the maximum learning rate 1 × 10^−4^, and a decayed period that comprised remain 90%, decreasing from the maximum learning rate 1 × 10^−4^ to 0. The AdamW optimizer was employed with its default settings to facilitate efficient optimization. The training batch size was set to 32, with gradient accumulation occurring every eight batches to manage memory effectively while achieving more stable gradient estimates. To speed up the training process, PyTorch Lightning (v2.0.4) (https://lightning.ai/pytorch-lightning) was implemented, enabling distributed computation across multiple GPUs. This framework significantly enhanced the training efficiency by parallelizing computations. ChromBERT underwent a rigorous training stage, spanning 100 epochs, which was completed in approximately 23 days with four NVIDIA A800 80GB GPUs. This intensive training period was necessary to ensure that the model adequately learned the complex patterns in the training data, setting a foundation for the subsequent fine-tuning stage.

### Representations of ChromBERT

#### TRN embeddings

ChromBERT is designed to represent context-specific TRNs by modeling the combinatorial binding patterns of transcription regulators across diverse genomic contexts. For each genomic region *b*_*i*_, ChromBERT encodes it into TRN embeddings 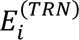 ∈ ℝ^*P*×*d*_*model*_^. These embeddings are generated through transformer blocks, which provide a flexible architecture for fine-tuning the model’s parameters to adapt to diverse downstream tasks. By adjusting attention mechanisms between transcription regulators during fine-tuning, ChromBERT captures the architectures of TRNs specific to the given biological contexts. These embeddings offer crucial insights and task-specific biological interpretations, underpinning the model’s utility in various transcription regulation-related applications.

Specifically, the 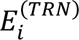 focused on the context-specific interactions of *P* (6,391) cistromes, which are pooled ChIP-seq profiles of *M* (1,073) regulators. To improve the model’s utility and interpretability, we transformed the cistrome-based TRN embeddings 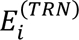 to regulator-based TRN embeddings 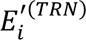 ∈ ℝ^*M*×*d_model_*^. This transformation involves characterizing the role of individual transcription regulators within the TRN. For regulators associated with multiple cistromes, we computed the average of all cistromes embeddings linked to a single regulator *r_m_* across different cell types at genomic bin *b*_*i*_, resulting in 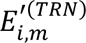 ∈ ℝ^*d*_*model*_^, which is defined as follows:

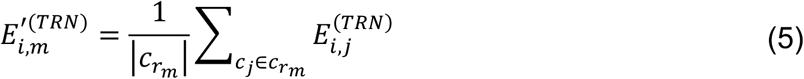

where *c*_*r*_*m*__ represent all cistromes associated with *r*_*m*_. These regulator-based embeddings were consistently applied in subsequent fine-tuning tasks and interpretability analysis by default.

#### Regulator embeddings

After transformation from cistrome-based TRN embedding to regulator-based TRN embedding, ChromBERT can output regulator embedding 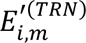 for each regulator *r*_*m*_ at genomic bin *b*_*i*_

#### Inference for functional collaborations of transcription regulators

In assessing the functional collaborations of transcription regulators, we inferred their interactions based on the similarity of their regulator embeddings within TRN presentations, derived from either a pre-trained model or a fine-tuned model adapting to specific downstream tasks. Specifically, for a given genomic region *b*_*i*_, we computed the embedding similarity between pairs of regulators, denoted as *r_j_* and *r*_*k*_. This was quantified using the cosine similarity formula:

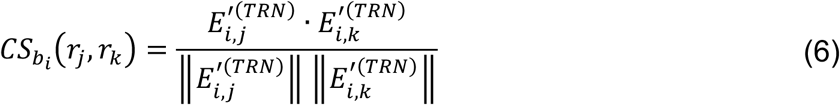

where 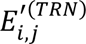 and 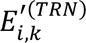 represents regulator embeddings for *r*_j_ and *r*_*k*_ at bin *b*_*i*_ from the TRN embeddings 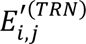. Furthermore, for functional collaborations across a specific set of bins, we averaged the regulator embeddings across all bin in the set. The embedding similarity was then calculated in a similar manner, enabling a broader understanding of regulator interactions across multiple genomic regions.

### In silico perturbation and omission

#### In silico perturbation

ChromBERT takes the genome-wide binding events of around one thousand transcription regulators as input, which can be in silico perturbed for analysis exploring the impact of binding events of individual regulator on the model output. This involved modifying the binding events of cistromes associated with each transcription regulator *r*_*m*_ across the genome. Specifically, we transformed the original input *X*_*i,j*_ into an in silico knockout input 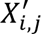, where converted all categories with positive or “slightly negative” binding status to “strongly negative” category. For each perturbed regulator *r*_*m*_, this in silico perturbation process was applied to all associated cistromes *c*_j_ ∈ *c*_*rm*_.

#### Omission of cistromes

Generally, ChromBERT processes binding status vector *X*_*i*_ = [*X*_*i*,1_, *X*_*i*,2_, ⋯, *X*_*i*,*P*_](*P* = 6,391). However, in certain analyses, highly dominant cistromes may overshadow the contributions of cofactors related to the learning objectives. For example, the presence of H3K27me3 is strongly correlated with EZH2’s classical role in gene expression repression. This dominance often leads to the exclusion of other regulators during fine-tuning tasks of classifying classical versus non-classical EZH2 binding sites. As a result, omitting H3K27me3 cistromes can enhance the identification of other key regulators. Formally, we yield a new binding status vector 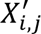 = [*X*_*i*,1_, *X*_*i*,2_, ⋯, *X*_*i*,*P*_] where *c*_j_ ∉ *c*_*r_m_*_ by omitting all cistrome related with *r*_*m*_.

### Fine-tuning of ChromBERT

#### Basic principle of fine-tuning

The fine-tuning of ChromBERT was conducted using a selective training strategy to effectively leverage its pre-trained knowledge. Initially, ChromBERT was loaded with pre-trained weights to retain the general representations learned during pre-training. To preserve this foundational knowledge, the early layers of the encoder were frozen, while the final layers of the encoder and all layers of the task-specific decoder were made trainable. This approach ensures that ChromBERT retains its pre-trained knowledge while adapting effectively to the specific requirements of diverse downstream tasks. By selectively adjusting the model, ChromBERT can adapt its TRN embeddings to align with task-specific and cell-type-specific objectives. This capability not only enables the model to perform well across a range of downstream tasks but also facilitates the interpretation of the roles and functional collaborations of transcription regulators in cell-type-specific transcription regulation.

#### Fine-tuning for cell-type-specific regulatory effects

ChromBERT was fine-tuned to model regulatory effects specific to cell types by learning genomic-context-specific labels across diverse genomic regions within each cell type. These labels correspond to the regulatory effects of specific genomic regions (e.g., active or inactive enhancers) in the given cell type. Generally, ChromBERT has adopted a universal design of decoder header architecture for these, eliminating the need for task-specific configurations. This design includes a convolutional layer followed by a multi-layer perceptron (MLP), which efficiently transforms TRN embeddings into the requisite outputs. The output can be a single probability for binary classification tasks or a single value for regression tasks, depending on the needs of the downstream tasks. Notably, this design also supports tasks with inputs subjected to in silico perturbation or omission. Further details on this kind of downstream tasks were described in the following downstream tasks section, and the frozen number of transformer blocks, loss functions and training hyperparameters were summarized in Supplementary Table S3.

#### Identification of key regulators in cell-type-specific regulatory effects

After fine-tuning ChromBERT to model regulatory effects in specific cell types, we analyzed the task-adapted TRN embeddings to identify key transcription regulators driving these cell-type-specific regulatory effects. Transformer-based deep language models, such as BERT, are particularly well-suited for this purpose due to their ability to generate contextualized embeddings that capture the context-dependent roles of individual elements^15,37^. ChromBERT leverages this capability to generate regulator embeddings that vary across genomic contexts (e.g., 1-kb genomic bins), allowing for a detailed analysis of transcription regulators’ roles within diverse genomic regions.

To identify key regulators, genomic regions are classified into two groups characterized by distinct regulatory effects. For each transcription regulator, the average embedding is computed across the regions in each group using the formula:

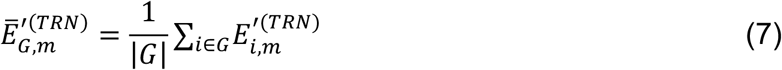

where *G* represent the set of 1-kb genomic bins and 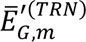 denotes the average embedding for the regulator *r*_*m*_ across the set of loci *G* derived from the fine-tuned model. Then, we evaluate the distance between embeddings of the two groups of loci for each transcription regulator as follows:

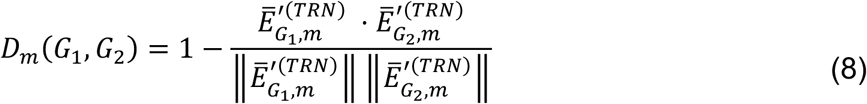

A greater distance *D*_*m*_ indicates higher contextual variability for the regulator, signifying its functional importance in distinguishing the regulatory effects between two groups. Transcription regulators with the highest embedding distances are identified as key regulators of context-specific regulatory effects within each task. This approach highlights the strength of ChromBERT, and other BERT-based architectures, in capturing the nuanced, context-dependent roles of transcription regulators. By interpreting regulator embeddings across functionally distinct loci, ChromBERT enables the identification of key regulators underlying the given genomic diversity. This provides valuable insights into transcriptional regulation within cell-type-specific settings, particularly in the absence of extensive experimental data for individual regulators.

#### Fine-tuning for regulatory dynamics between cell types

In our study, we utilized ChromBERT to understand transcription regulatory dynamics in single cellular heterogeneity and during cell state transitions. Specifically, the model was fine-tuned to predict genome-wide changes in transcriptome and chromatin accessibility between pairwise cell types. The fine-tuning process for chromatin accessibility followed the same objectives as those used for determining cell-type-specific regulatory effects. However, the objectives for transcriptome changes differed due to the potential uncertainty in the influence of regions adjacent to transcription start sites (TSSs) on gene expression. For each gene, we included both upstream and downstream regions relative to the TSS to capture a comprehensive genomic context. We selected four genomic bins on either side of the TSS, as well as the bin containing the TSS itself, denoted as [*b*_*i*–4_, *b*_*i*–3_, *b*_*i*–2,_*b*_*i*–1,_*b*_*i*,_*b*_*i*+1_, *b*_*i*+2_, *b*_*i*+3,_*b*_*i*+4_]. To derive the most informative TRN embeddings from these regions, we then employed a max pooling strategy. This method ensures that the most significant signals across these genomic bins are captured, enhancing the accuracy of our predictions regarding gene expression regulation. A universal decoder architecture was followed to transform the TRN embeddings to the requisite output. Further details on this kind of downstream tasks were also described in the following downstream tasks section, and Supplementary Table S3.

#### Identification of key regulators in regulatory dynamics between cell types

Following fine-tuning for regulatory dynamics, ChromBERT identifies key transcription regulators critical to changes in regulatory effects between cell types. This process parallels the identification of key regulators in cell-type-specific regulatory effects, with the difference being that the embedding distances are calculated between groups of regions categorized as having changed or unchanged regulatory effects between cell types. This enables ChromBERT to pinpoint regulators driving transcriptional dynamics across cell types, providing insights into the mechanisms underlying cellular transitions.

#### Fine-tuning with prompts for additional information incorporation

Due to the mixed cell type pooling operation and reliance on cistromes involved in pre-training, the encoder of ChromBERT exhibits a limited capacity to generate variable TRN embeddings in response to specific inputs. In this study, we employed a prompt-enhanced fine-tuning approach, integrating TRN representations with additional relevant information, including cell-type-specific data or genetic variations. This approach successfully addresses these limitations and broadens the model’s applicability across diverse objectives, including cistromes imputation and the identification of causal eQTLs. By annotating cistromes, cell-type-specific chromatin accessibility representation derived from pre-trained ChromBERT embedding can be utilized as cell-type-specific information for cistromes imputation. Additionally, transcriptome embeddings from scGPT were also employed as cell-type-specific information in this study. Genetic variation information we used was sourced from DNABERT-2 for identification of causal eQTLs. Vectors containing additional information were denoted as prompts in this study. To utilize the TRN representations and prompts, we concatenated each TRN embeddings 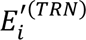 with respective prompts 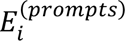. In this concatenation, we performed a mean pooling on sequence length dimension to convert the TRN embeddings into a 768-dimensional vector for convenient concatenation with prompts. This concatenation produced a prompt-enhanced embeddings 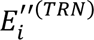, which was subsequently processed by an MLP decoder to generate the necessary outputs. Further details on this kind of downstream tasks were described in the following downstream tasks section.

### Downstream tasks

#### Hi-C imputation

We employ pre-trained TRN embeddings to enhance Hi-C contact map imputation, effectively capturing spatial chromatin regulation. We employed a CNN model utilizing three input channels: a O/E normalized Hi-C contact map, a pairwise genomic distance map expressed as (1 + *Dg*)^−0.75^, where *Dg* represents genomic distance, and a pairwise TRN embedding distance map defined by cosine similarity. The input window is set at 1000 kb with a 1-kb resolution and moves diagonally in 200-kb steps. Hi-C contact maps were initially processed at 5-kb resolution and then linearly interpolated to a 1-kb resolution. We used an O/E normalized Micro-C contact map as the ground truth, dividing the chromosomes into three sets for training, validation, and testing. The training set includes chromosomes 1, 4, 7, 10, 13, 17, and 18; the validation set consists of chromosomes 3, 6, 9, 12, 16, 19, 20, and X; and the test set contains chromosomes 2, 5, 8, 11, 14, 15, 21, and 22. Specially, 1-kb bins excluded during ChromBERT’s pre-training were assumed to lack TF binding, generating embeddings with an all-zero binding status vector. To minimize potential data bias, we adjusted the cosine similarity values for contacts involving these excluded regions by substituting them with the mean cosine similarity value of contacts at the same distance.

#### Cistrome imputation using cell-type-specific prompts (fine-tuning with prompts)

The current matrix representing cell types and transcription regulators is highly sparse due to the lack of available cistromes for many transcription regulators across diverse cell types. To address this issue, our method integrates cell-type-specific prompts based on chromatin accessibility into the model to impute the matrix by predicting new cistromes, leveraging chromatin accessibility’s established role in predicting transcription regulator binding statuses. Alternatively, cell-type-specific prompts based on transcriptomic profiles can be incorporated, as these profiles are also linked to transcription regulator binding statuses and are abundantly available. Furthermore, our approach incorporates a regulator prompt to differentiate among various transcription regulators, facilitating a universal generative model architecture capable of generating cistromes for distinct transcription regulators and performing binary classification to determine their presence or absence. These prompts improve the model’s ability to adapt its predictions to specific cell types.

- Chromatin accessibility prompt: We leveraged DNase-seq data extensively available for various cell types in the Cistrome-Human-6K. For each cell type, ChromBERT’s pre-trained embeddings specifically for DNase-seq data were used, denoted as 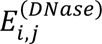 ∈ ℝ^*d*_*model*_^, where *i* represents the *i*-th bin and *j* indicates the position index of DNase-seq data within the Cistrome-Human-6K dataset for the given cell type.
- Transcriptome prompt: Transcriptome prompts were derived from cell embeddings of scGPT^6^ following zero-shot learning on transcriptomic profiles, either from single-cell or bulk cell data, specific to the cell type, denoted as *E*^(*scGPT*)^ ∈ ℝ^*dscGPT*^, where *d*_𝑠*cGPT*_ = 512. For single-cell RNA-seq data, cells covering fewer than 200 or more than 7,000 genes were filtered out, and the sum of read counts in each cell were normalized to 1 × 10^4^ followed by a log1p transformation using Scanpy (v1.9.5)^61^. scGPT then processed these profiles, only considering genes with non-zero expression levels for the computation of attention. For bulk cell RNA-seq data, we utilized the same normalization and transformation procedure and used top 7,000 highly-expressed genes to obtain cell embeddings.
- The regulator prompt was pre-trained regulator embedding 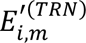 for given regulator *r*_*m*_ at the *i*-th 1-kb bin.

Then for this fine-tuning task, the prompts can be:

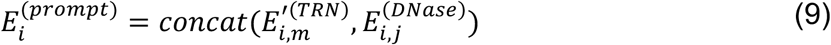

or

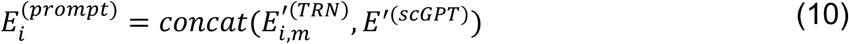

based on the choice of cell-type-specific information representation, where *E*^′(*scGPT*)^represents transformed *E*^(*scGPT*)^by a linear layer to match the hidden dimensions for concatenation. Remarkably, *E*^′(*scGPT*)^is cell representations and is consistent across all bins in a certain cell type.

To fine-tune the model for cistrome prediction across various cell types and transcription regulators using chromatin accessibility as cell-type-specific prompts, the model was trained with DNase-seq data from 22 cell types and utilized 104 cistromes corresponding to 29 transcription regulators within these cell types (Supplementary Table S4). For the model fine-tuned with transcriptomic prompts as cell-type-specific indicators, training was conducted using scGPT cell prompts derived from the RNA-seq dataset, encompassing 20 cell types and 102 cistromes for 29 transcription regulators within these cell types (Supplementary Table S4). After the training, the fine-tuned model based on chromatin accessibility prompts or transcriptomic prompts were both applied to perform prediction in bulk cell types for transcription regulators.

And the fine-tuned model based on transcriptome prompts was further applied to human PBMC 10x single-cell multi-omics dataset. Paired single-cell RNA-seq and ATAC-seq data were collected from the 10x Genomics multi-omics repository (https://www.10xgenomics.com/datasets/pbmc-from-a-healthy-donor-granulocytes-removed-through-cell-sorting-10-k-1-standard-2-0-0). Initial data processing and quality control for single-cell RNA-seq data were performed according to protocols in muon-tutorials (https://muon-tutorials.readthedocs.io/en/latest/single-cell-rna-atac/pbmc10k/1-Gene-Expression-Processing.html). We then processed to cluster cells and annotated different cell clusters based on marker gene expression profiles as describe in the same tutorial. Similar data processing and quality control steps were performed for the single-cell ATAC-seq data according to the muon-tutorials (https://muon-tutorials.readthedocs.io/en/latest/single-cell-rna-atac/pbmc10k/2-Chromatin-Accessibility-Processing.html). After these initial quality checks, we further refined the dataset by excluding single-cell ATAC-seq reads detected in fewer than 10 cells and cells with fewer than 1,000 detected genes in single-cell RNA-seq data. With this workflow, we obtained a final 9,493 cells from an initial pool of 11,898 cells. After data processing and cell clustering, we applied fine-tuned ChromBERT with single-cell RNA-seq embeddings as transcriptome prompts to perform prediction in single cells for transcription regulators. Specifically, in this prediction section, we removed all cell clusters related to T cells as we observed that scGPT cell embeddings could not distinguish T cells from other cell clusters well.

#### Fine-mapping of eQTLs with DNA sequence prompt (fine-tuning with prompts)

To fine-tune ChromBERT for fine-mapping of eQTLs with DNA sequence prompt, we concatenated mean pooled TRN embeddings with DNA sequence prompt to be fed into an MLP decoder to classify the causal and non-causal eQTLs. For each eQTL, we obtained DNA sequence embeddings with DNABERT-2 for a 1-kb window (from upstream 500 bp to downstream 499 bp of the coordinate of the eQTL). The variant DNA sequence embeddings 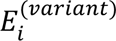 were used as the DNA sequence prompt.

For this classification task, we collected eQTLs from the EBI eQTL catalogue (https://www.ebi.ac.uk/eqtl) which utilized the SuSiE protocol as described in the Enformer study^7,62^. We systematically analyzed eQTLs across 49 different tissues. For each tissue, variants demonstrating a high likelihood of influence on gene expression, indicated by a posterior inclusion probability (PIP) > 0.9, were categorized as causal variants. Conversely, for each identified causal variant, we sought a corresponding non-causal variant. This non-causal variant was chosen from the set with a PIP < 0.01 but an absolute *Z*-score > 4 tested for the same gene, wherever available. In cases where a matched gene variant was not available, we instead selected a random variant from the genome-wide locus with a PIP < 0.01 and an absolute *Z*-score > 6, to serve as the non-causal counterpart. Following the approach used in Enformer study^7^, we fine-tuned the model in each tissue by conducting 10 iterations of cross validation, maintaining a training/testing ratio of 6:4.

#### Prediction for genome-wide STARR-seq signals (fine-tuning for cell-type-specific regulatory effects)

For this task, we utilized STARR-seq data from wild type HCT116 cells available in GSE156740^44^. This involved downloading both the STARR-seq signals and a reference set of enhancers, which are based on the human genome build hg19. To align this with ChromBERT, which uses hg38, we converted hg38 1-kb bins to hg19 using the liftOver tool (https://hgdownload.soe.ucsc.edu/admin/exe/linux.x86_64/liftOver). We then associated the human hg38 1-kb bins with the reference STARR-seq enhancers and processed the STARR-seq signals using a log_2_-transformed fold changes with the pseudo count one. The model was trained specifically on regions that included these reference enhancers and additional open chromatin regions identified by DNase-seq data in HCT116 cells (ENCFF240LRP). This training approach was designed to enhance robustness of our model for variance of STARR-seq signals. Importantly, we excluded all cistromes related to histone modifications or chromatin accessibility to focus specifically on the influence of transcription factors on enhancer activity.

#### Prediction for perturbation effect on STARR-seq signals (fine-tuning for cell-type-specific regulator effects)

For predicting the perturbation effects on STARR-seq signals, we adapted ChromBERT to predict changes in control and 3-indoleacetic acid (IAA)-treated HCT116 cells. IAA treatment is known to rapidly deplete tagged transcription factors^44^. We used the original reference cistromes for control cell predictions and applied in silico perturbations for binding status of depleted factors in treated cells (see the in silico perturbation section above). Training for this in silico perturbation model included STARR-seq signals in both control (-IAA) and treated (+IAA) conditions for three transcription factors: BRD2, CDK7, and EP300/CREBBP. Unlike the model for wild type signals, this in silico perturbation model was trained using reference STARR-seq enhancers that show significant variance in response to treatment.

#### Classification of EZH2’s classical and non-classical sites (fine-tuning for cell-type-specific regulatory effects)

For this classification task, we used EZH2 (GSE29611^2^) and H3K27me3 (GSE61176) ChIP-seq data from human embryonic stem cells. We labeled bins overlapping with both EZH2 and H3K27me3 as positive and those only overlapping with EZH2 as negative to create a clear binary classification dataset. As H3K27me3 was included in the pre-trained dataset and was proposed to have dominant effect on the model output, we omitted cistromes associated with H3K27me3 from the input reference cistromes.

#### Prediction for genome-wide changes in transcriptome or chromatin accessibility (fine-tuning for regulatory dynamics between cell types)

In this task, we fine-tuned ChromBERT to predict genome-wide changes in transcriptome or chromatin accessibility in cellular heterogeneity and during transdifferentiation. Specifically, we quantified genome-wide changes in transcriptome with fold changes of gene expression levels, and quantified genome-wide changes in chromatin accessibility through fold changes of DNase-seq or ATAC-seq signals.

After data processing and cell clustering for human PBMC 10x single-cell multi-omics dataset mentioned in the above sections in Methods. We specifically assigned cell clusters into five major cell types for this prediction section, including T cells, B cells, monocytes, dendritic cells (DCs) and natural killer (NK) cells. This assignment helps us to collected well-known key regulators for each cell type from the public studies. For each cell type, single-cell RNA-seq data were aggregated into pseudo-bulk counts and normalized using a factor of 1 × 10^6^. Single-cell ATAC-seq data were also compiled into pseudo-bulk reads and analyzed for peak calling using pycisTopic (v1.0.3)^56^, and these reads were normalized to a count of 1 × 10^6^. Then we fine-tuned ChromBERT to predict changes in transcriptome or chromatin accessibility between cell types in a pairwise manner. For transcriptome profiles, we calculated changes using differences of log1p-transformed pseudo-bulk expression level across all genes. For chromatin accessibility, changes were calculated by log_2_-transformed fold changes in pseudo-bulk reads density at merged peaks and background regions (spanning 10 kb upstream and downstream around all TSSs), with a pseudo count one in fold change calculation.

For transcriptome changes in transdifferentiation, we collected transcriptome profiles derived from CAGE-seq in seven cell types available in the FANTOM5 database^63^, including fibroblasts, induced pluripotent stem cells (iPSCs), myoblasts, hepatocytes, macrophages, cardiac myocytes, and B cells. We focused on five transdifferentiation processes: fibroblast to iPSCs, fibroblast to myoblast, fibroblast to hepatocyte, fibroblast to cardiac myocyte, and B cells to macrophage. Bulk-cell transcriptome changes across different cell states were quantified with differences of log1p-transformed gene expression level (TPM) across all genes.

For chromatin accessibility changes in transdifferentiation, we utilized DNase-seq in fibroblast (ENCFF184KAM), iPSCs (ENCFF540VPT), myoblast (ENCFF647RNC), cardiac myocyte (ENCFF389SOW), B cells (ENCFF650BNV) and macrophage (ENCFF580ICE), and ATAC-seq in fibroblast and hepatocyte (GSE179011^64^, GSE185358^65^). Peak calling and reads density normalization were performed in the approaches similar to those used in our single-cell ATAC-seq pseudo-bulk processing. And bulk-cell chromatin accessibility changes were also quantified by log_2_-transformed fold changes in reads density at merged peaks and background regions (spanning 10 kb upstream and downstream around all TSSs), with a pseudo count one in fold change calculation.

We performed interpretation analysis to identify key regulators in cellular heterogeneity and during cell state transitions. In the realm of cellular heterogeneity, we categorized genes and loci based on elevated or stable expression and chromatin accessibility. Specifically, genes exhibiting a log1p-transformed expression difference greater than 1 were classified as having increased expression, while those with a difference ranging from −0.5 to 0.5 were considered stable. In chromatin accessibility analysis using ATAC-seq, regions with a log_2_-transformed increase greater than 2 were marked as having increased chromatin accessibility. After excluding regions that had no coverage, we selected the top 40,000 regions showing the minimal absolute fold changes as having unchanged chromatin accessibility. The method for defining genes and regions with increased or unchanged expression or accessibility during cell state transitions was analogous to that used for analyzing cellular heterogeneity.

Each analysis between two single-cell pseudo-bulk cell types in cellular heterogeneity or two cell states in a transdifferentiation process involved ranking transcription factors based on their embedding distance in two set of loci for transcriptome or chromatin accessibility dynamics, respectively. Epigenetic modifications were not considered in this analysis. We averaged ranking scores from these two modalities to compute a composite ranking score for 991 transcription factors, identifying those most influential in driving cellular state transition.

In our analysis of cell state transition in each transdifferentiation process, the top 25 transcription factors were recognized as key regulators of this transdifferentiation process. For cellular heterogeneity, we computed an average ranking score for each regulator by averaging ranking scores for the given cell type relative to each other (defined as cell-type-specific ranking scores). This detailed evaluation helps us understand the importance of these key regulators in differentiating the given cell type from all other cell types and identify the top 25 key regulators for each cell type, 84 key regulators in total. And cell-type-specific ranking scores for all 84 key regulators help us to further assign the cell-type-specificity of identified key regulators, which were generally classified into four categories, B cells-specific, T cells/NK cells-specific, monocytes-specific and DCs-specific, as illustrated in Fig. 4b.

### Benchmarks

#### Benchmark for cistrome imputation

We assessed the performance of ChromBERT on the cistrome imputation task by comparing it with both Avocado and a baseline approach based on putative peaks and chromatin accessibility. For the Avocado comparison, we used imputed tracks from ENCODE (ENCSR481OSA) and restricted our analysis to cistromes withheld from training. For benchmarking, imputed signals derived from BigWig files were aggregated into 1-kb bins.

The baseline method was established by integrating putative peaks with chromatin accessibility. For each test cistrome, putative peaks were identified as follows. If at least five cistromes (excluding the test cistrome) were present in our Cistrome-Human-6K dataset, related peaks were collected and merged into 1-kb bins to define putative peaks. If fewer than five were available, putative peaks were instead derived from 1-kb bins based on the corresponding regulator’s motif. The prediction probability in the benchmark analysis was then computed by multiplying the average DNase-seq signal (aggregated over 1-kb bins) by a binary indicator (0 or 1) denoting the presence of putative peaks.

We evaluated ChromBERT’s performance to predict transcription regulator binding sites that are specific to a particular cell type, even when chromatin accessibility is similar across cell types. To define these sites, we performed pairwise comparisons between paired cistromes from different cell types but associated with the same transcription regulator. A binding site was classified as cell-type-specific if it was present in one cell type but absent in the other, represented as 1-kb bins. For a site to be considered peak-present, it had to meet two criteria: (1) the signal intensity had to exceed 3; (2) the signal had to be at least threefold higher than in the peak-absent cell type. To control for potential differences in chromatin accessibility, we applied additional constraints: (1) the absolute log_2_-transformed fold change in DNase-seq signal between the two cell types had to be less than 0.5; (2) the DNase-seq signal in both cell types had to exceed 3, ensuring that both sites were located in accessible chromatin. ChIP-seq and DNase-seq signals were normalized to the genomic background based on read densities. To ensure statistical robustness, we included only cistrome pairs with at least 50 cell-type-specific binding events, while also requiring that the ratio of peak-present to peak-absent bins remained balanced, with a maximum difference of less than fivefold. ChromBERT’s performance to distinguish cell-type-specific binding events was assessed by comparing its predicted probability differences between paired cell types against the ground truth—defined by the actual presence or absence of transcription regulator binding based on ChIP-seq data.

#### Benchmark with XGBoost

The XGBoost model (implemented by Python package XGBoost, version 2.1.3) was utilized as a baseline to demonstrate the effectiveness of TRN embeddings generated by ChromBERT across various downstream tasks, using the same input as ChromBERT. While other tasks can directly utilize the XGBoost model, fine-tuning tasks with prompts require specific configurations, as direct training is not practical.

For cistrome imputation tasks based on DNase-seq prompts, the input vector is constructed by concatenating the following three components: a discrete signal token for the DNase-seq in the target cell type, serving as a chromatin accessibility prompt; a discrete signal token for cistromes associated with the target transcription regulator indicated by the most frequent one, functioning as the regulator prompt; and the cistromes binding status vector *X*_*i*_ = [*X*_*i*,1_, *X*_*i*,2_, ⋯, *X*_*i*,*P*_]. The training and validation of the XGBoost model was conducted on the same datasets used by ChromBERT, following an incremental learning strategy with the binary cross-entropy (BCE) loss function. The training dataset was divided into 356 randomly assigned blocks, processed sequentially. Initially, the first block permitted up to 50 rounds of boosting, implementing early stopping if no improvement was detected on the validation dataset for 10 consecutive rounds. For subsequent blocks, the model employed a number of boosting rounds equivalent to the number of trees in the prior block, maintaining the same early stopping methodology.

For the task of fine-mapping of eQTLs, the input vector fed into XGBoost is a concatenated vector comprising variant DNA sequence embeddings 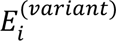 and the cistromes binding status vector *X*_*i*_. All other parameters use the default settings.

#### Benchmark with Enformer and DNABERT-2

In our study, we also compared the performance of ChromBERT with two advanced deep language models, Enformer^7^ and DNABERT-2^8^, in a subset of our downstream tasks which can also be accomplished by the two models based on DNA sequences (genome build hg38). These models, which generate contextual representations for input DNA sequences, were fine-tuned using the established method in the original studies with slight modification to suit specific requirements of the tasks. For DNABERT-2, we implemented a fine-tuning protocol available from the study (https://github.com/MAGICS-LAB/DNABERT_2/blob/main/finetune/train.py). In the case of Enformer, we used a random forest approach, either as a classifier or a regressor, to perform predictions on downsteam tasks based on the embeddings generated from DNA sequences. The random forest classifier and regressor were implemented using the package scikit-learn (v1.3.0)^66^, with the “max_features” parameter set to “log2”. The number of frozen transformer layers, loss functions and training hyperparameters for DNABERT-2 and Enformer were summarized in Supplementary Table S3.

Both models were trained using the same dataset as ChromBERT. DNABERT-2 processed the entire DNA sequences in 1-kb bins, similar to ChromBERT. If ChromBERT was set to consider multiple adjacent bins (e.g., *n* = 9 bins for changes in the cellular transcriptome), DNABERT-2 followed the same approach and then applied a max pooling strategy to the embeddings. Enformer, designed to handle long-range DNA sequences, required inputs of 196,608 bp. For this purpose, we used one-hot-encoded DNA sequences centered around the 1-kb bins used in ChromBERT. These inputs were then transformed into 896 sequence vectors by Enformer, corresponding to 114,689 bp in 128-bp resolution. To match the lengths of input regions from different models, we selected the central 8 out of 896 sequence vectors from Enformer’s output, corresponding to 1,024 bp, as Enformer’s embeddings for the given 1-kb bin. For these 8 sequence vectors, we performed a mean pooling strategy to generate an averaged sequence vector for the following random forest.

For tasks where ChromBERT used multiple adjacent bins (e.g., *n* = 9 bins for changes in the cellular transcriptome), we selected the central 70 out of 896 sequence vectors from Enformer’s output, corresponding to 8,960 bp, and applied a similar mean pooling strategy.

#### Gene ontology enrichment analysis

We conducted Gene Ontology (GO) enrichment analysis using the DAVID online tool (https://david.ncifcrf.gov/tools.jsp)^67^. This analysis assessed the enrichment of human gene ontology biological process (BP) terms.

#### Motif scan

Motif scans were performed using FIMO (v5.0.5)^69^ against the JASPAR core 2024 vertebrates database^70^ with the following parameters “--max-stored-scores 1000000”. Motifs with p-value ≤ 1 × 10^-5^ were used for the following analysis.

## Data availability

Cistrome-Human-6K dataset is available on the Hugging Face Dataset Hub (https://huggingface.co/datasets/TongjiZhanglab/chrombert). The blacklist of genomic regions was obtained from the ENCODE project (https://www.encodeproject.org) under accession code ENCFF356LFX. ChIP-seq data used for analysis in Supplementary Fig.S2h was obtained from GEO accession code GSE62562 (H3K27me3)^68^ and GSE39912 (H3K4me3)^69^. The Hi-C dataset used for the analysis of Supplementary Fig. S2c was obtained from GEO accession GSE63525^70^. The Micro-C and Hi-C data in hESC cells (4DNES21D8SP8, 4DNES2M5JIGV) and HFF cells (4DNESWST3UBH, 4DNES2R6PUEK^71^) used for Hi-C imputation in Supplementary Fig. S2d-e were obtained from 4DN consortium (https://data.4dnucleome.org/). The ChIP-seq of BRD4 (GSE89128^72^, GSE99178^73^) and DNase-seq data (GSE26328^74^, GSE25344^75^) in A549 and K562 cells used in Fig. 2c-d are obtained from GEO. The ChIP-seq of EP300 (GSE51176^76^, GSE59681^77^) and DNase-seq data (GSE50610^78^, GSM723024^79^) in HCT116 and IMR-90 cells used in Fig. S3c are obtained from GEO. The data used in Fig 2b, Fig 2e-g and Supplementary Fig. S4a-b were obtained from ENCODE and GEO, whose file accession codes were available in Supplementary Table S2. eQTLs are available from the EBI eQTL catalogue (https://www.ebi.ac.uk/eqtl). ChIP-seq data of YY1 and DNase-seq data used for the analysis in Fig. 3d was obtained from GEO accession GSE32465^2^ and GSE18927^79^. The DNase-seq in HCT116 used to define open regions were downloaded from (https://www.encodeproject.org) with file accession code ENCFF240LRP and the list of enhancer regions for analysis in Fig.3e-h is available from the supplementary material of the previous study^44^. The STARR-seq data used for analysis in Fig.3e-h were downloaded from accession code GSE156740^44^. ChIP-seq data of EP300 and BRD4 used in Fig. 3g were from accession codes GSE51176^76^ and GSE57628^80^. ChIP-seq data in hESC used for analysis in Fig. 3i was obtained from GEO accession codes GSE61176^81^ (H3K27me3) and GSE29611^2^ (EZH2). The RNA-seq data used as transcriptome prompt of ChromBERT in the cistrome imputation task were obtained from ENCODE and the file accession codes were listed in Supplementary Table S4. The single-cell RNA-seq and ATAC-seq data used in cistrome imputation task of Supplementary Fig. S4 was obtained from PBMC 10x single-cell multi-omics dataset (https://www.10xgenomics.com/datasets/pbmc-from-a-healthy-donor-granulocytes-removed-through-cell-sorting-10-k-1-standard-2-0-0). The DNase-seq data used for modeling the chromatin accessibility changes in transdifferentiation was obtained from (https://www.encodeproject.org) with file accession codes ENCFF184KAM (fibroblast), ENCFF540VPT (iPSC), ENCFF647RNC (myoblast), ENCFF389SOW (cardiac myocyte), ENCFF650BNV (B cell) and ENCFF580ICE (macrophage). The ATAC-seq data used for modeling the chromatin accessibility changes in transdifferentiation was downloaded from accession codes GSE179011^64^ and GSE185358^82^. The CAGE-seq data used for modeling the transcriptome changes in transdifferentiation was obtained from (https://fantom.gsc.riken.jp/5/datafiles/reprocessed/hg38_latest/extra/CAGE_peaks_expression/hg38_fair+new_CAGE_peaks_phase1and2_tpm.osc.txt.gz).

## Code availability

The pre-trained model, reference datasets and all source codes of ChromBERT are available on the GitHub repository (https://github.com/TongjiZhanglab/ChromBERT).

## Acknowledgments

We would like to thank Prof. Rui Jiang for comments on designing the research. This work was supported by the National Natural Science Foundation of China (32325012 (Y(Yong).Z.), 32030022 (Y(Yong).Z.), 32488101 (Y(Yong).Z.), 32400522 (Z.Y.)), the National Key Research and Development Program of China (2021YFA1302500 (Y(Yong).Z.)), the Science and Technology Commission of Shanghai Municipality (23JS1401200 (Y(Yong).Z.)), the Postdoctoral Innovation Talents Support Program (BX20230265 (Z.Y.)), the China Postdoctoral Science Foundation (2022M722423 (Z.Y.)) and the GHfund C (202302033256 (Z.Y.)).

## Author Contributions

Y(Yong).Z. conceived the research, Y(Yong).Z. and Z.Y. designed the research. Z.Y. and D.Y. developed the model, Q.C. and Y(Yuxuan).Z. performed computational analysis with the help of Y.W., Z.L. and C.W., Z.Y., D.Y., Q.C., Y(Yuxuan).Z. and Y(Yong).Z. wrote the manuscript.

## Competing interests declaration

The authors declare they have no competing interests.

## Notes

### Competing Interest Statement

The authors have declared no competing interest.

### Summary of Updates

The original author list present in bioRxiv contained inaccuracies. In this revision, we have corrected the author list to accurately reflect the contributions of all authors.

## References

1. Gerstein, M.B. et al. Architecture of the human regulatory network derived from ENCODE data. Nature 489, 91–100 (2012).

2. Consortium, E.P. An integrated encyclopedia of DNA elements in the human genome. Nature 489, 57–74 (2012).

3. Stampfel, G. et al. Transcriptional regulators form diverse groups with context-dependent regulatory functions. Nature 528, 147–51 (2015).

4. Consortium, E.P. et al. Expanded encyclopaedias of DNA elements in the human and mouse genomes. Nature 583, 699–710 (2020).

5. Theodoris, C.V. et al. Transfer learning enables predictions in network biology. Nature 618, 616–624 (2023).

6. Cui, H. et al. scGPT: toward building a foundation model for single-cell multi-omics using generative AI. Nat Methods (2024).

7. Avsec, Z. et al. Effective gene expression prediction from sequence by integrating long-range interactions. Nat Methods 18, 1196–1203 (2021).

8. Zhou, Z. et al. Dnabert-2: Efficient foundation model and benchmark for multi-species genome. arXiv preprint arXiv:2306.15006 (2023).

9. Hao, M. et al. Large-scale foundation model on single-cell transcriptomics. Nat Methods 21, 1481–1491 (2024).

10. Fu, X. et al. A foundation model of transcription across human cell types. Nature (2025).

11. Nguyen, E. et al. Sequence modeling and design from molecular to genome scale with Evo. Science 386, eado9336 (2024).

12. Dalla-Torre, H. et al. Nucleotide Transformer: building and evaluating robust foundation models for human genomics. Nat Methods (2024).

13. Gururangan, S., et al. Don’t stop pretraining: Adapt language models to domains and tasks. arXiv preprint arXiv:2004.10964 (2020).

14. Chen, V. et al. Applying interpretable machine learning in computational biology-pitfalls, recommendations and opportunities for new developments. Nat Methods 21, 1454–1461 (2024).

15. Ethayarajh, K. How contextual are contextualized word representations? Comparing the geometry of BERT, ELMo, and GPT-2 embeddings. arXiv preprint arXiv:1909.00512 (2019).

16. Yu, Z. & Zhang, Y. Foundation model for comprehensive transcriptional regulation analysis. Natl Sci Rev 11, nwae355 (2024).

17. Kamimoto, K. et al. Dissecting cell identity via network inference and in silico gene perturbation. Nature 614, 742–751 (2023).

18. Slattery, M. et al. Absence of a simple code: how transcription factors read the genome. Trends Biochem Sci 39, 381–99 (2014).

19. Yu, Z., Wang, Q., Zhu, G., Zhu, J. & Zhang, Y. Decoding the genomic landscape of chromatin-associated biomolecular condensates. bioRxiv, 2023.08. 23.554542 (2023).

20. Vaswani, A. et al. Attention is all you need. Advances in neural information processing systems 30(2017).

21. Devlin, J., Chang, M.-W., Lee, K. & Toutanova, K. Bert: Pre-training of deep bidirectional transformers for language understanding. arXiv preprint arXiv:1810.04805 (2018).

22. Taing, L. et al. Cistrome Data Browser: integrated search, analysis and visualization of chromatin data. Nucleic Acids Res 52, D61–D66 (2024).

23. Liu, T., et al. Cistrome: an integrative platform for transcriptional regulation studies. Genome Biol 12, R83 (2011).

24. Vu, H. & Ernst, J. Universal annotation of the human genome through integration of over a thousand epigenomic datasets. Genome Biol 23, 9 (2022).

25. Bonev, B. & Cavalli, G. Organization and function of the 3D genome. Nat Rev Genet 17, 772 (2016).

26. Huttlin, E.L. et al. Dual proteome-scale networks reveal cell-specific remodeling of the human interactome. Cell 184, 3022–3040 e28 (2021).

27. Liberzon, A. et al. The Molecular Signatures Database (MSigDB) hallmark gene set collection. Cell Syst 1, 417–425 (2015).

28. Wang, H. et al. Role of histone H2A ubiquitination in Polycomb silencing. Nature 431, 873–8 (2004).

29. Hu, S. et al. ncHMR detector: a computational framework to systematically reveal non-classical functions of histone modification regulators. Genome Biol 21, 48 (2020).

30. Liu, P. et al. Pre-train, prompt, and predict: A systematic survey of prompting methods in natural language processing. ACM Computing Surveys 55, 1–35 (2023).

31. Kirillov, A., et al. Segment anything. arXiv preprint arXiv:2304.02643 (2023).

32. Chowdhery, A. et al. Palm: Scaling language modeling with pathways. Journal of Machine Learning Research 24, 1–113 (2023).

33. Wei, J., et al. Finetuned language models are zero-shot learners. arXiv preprint arXiv:2109.01652 (2021).

34. Schreiber, J., Durham, T., Bilmes, J. & Noble, W.S. Avocado: a multi-scale deep tensor factorization method learns a latent representation of the human epigenome. Genome Biol 21, 81 (2020).

35. Grosselin, K. et al. High-throughput single-cell ChIP-seq identifies heterogeneity of chromatin states in breast cancer. Nat Genet 51, 1060–1066 (2019).

36. Bartosovic, M., Kabbe, M. & Castelo-Branco, G. Single-cell CUT&Tag profiles histone modifications and transcription factors in complex tissues. Nat Biotechnol 39, 825–835 (2021).

37. Wiedemann, G., Remus, S., Chawla, A. & Biemann, C. Does BERT make any sense? Interpretable word sense disambiguation with contextualized embeddings. arXiv preprint arXiv:1909.10430 (2019).

38. Uffelmann, E. et al. Genome-wide association studies. Nature Reviews Methods Primers 1, 59 (2021).

39. Boyle, A.P. et al. High-resolution mapping and characterization of open chromatin across the genome. Cell 132, 311–22 (2008).

40. Song, L. & Crawford, G.E. DNase-seq: a high-resolution technique for mapping active gene regulatory elements across the genome from mammalian cells. Cold Spring Harb Protoc 2010, pdb prot5384 (2010).

41. Wang, Y. et al. SNP rs17079281 decreases lung cancer risk through creating an YY1-binding site to suppress DCBLD1 expression. Oncogene 39, 4092–4102 (2020).

42. Arnold, C.D. et al. Genome-wide quantitative enhancer activity maps identified by STARR-seq. Science 339, 1074–7 (2013).

43. Krebs, A.R., Karmodiya, K., Lindahl-Allen, M., Struhl, K. & Tora, L. SAGA and ATAC histone acetyl transferase complexes regulate distinct sets of genes and ATAC defines a class of p300-independent enhancers. Mol Cell 44, 410–423 (2011).

44. Neumayr, C. et al. Differential cofactor dependencies define distinct types of human enhancers. Nature 606, 406–413 (2022).

45. Margueron, R. & Reinberg, D. The Polycomb complex PRC2 and its mark in life. Nature 469, 343–9 (2011).

46. Xu, K. et al. EZH2 oncogenic activity in castration-resistant prostate cancer cells is Polycomb-independent. Science 338, 1465–9 (2012).

47. Xu, H. et al. Integrative analysis reveals the transcriptional collaboration between EZH2 and E2F1 in the regulation of cancer-related gene expression. Molecular Cancer Research 14, 163–172 (2016).

48. Kim, E. et al. Phosphorylation of EZH2 activates STAT3 signaling via STAT3 methylation and promotes tumorigenicity of glioblastoma stem-like cells. Cancer Cell 23, 839–52 (2013).

49. Laidlaw, B.J. & Cyster, J.G. Transcriptional regulation of memory B cell differentiation. Nat Rev Immunol 21, 209–220 (2021).

50. Fontenot, J.D. et al. Regulatory T cell lineage specification by the forkhead transcription factor foxp3. Immunity 22, 329–41 (2005).

51. Fang, D. et al. Differential regulation of transcription factor T-bet induction during NK cell development and T helper-1 cell differentiation. Immunity 55, 639–655 e7 (2022).

52. Chopin, M. et al. Transcription Factor PU.1 Promotes Conventional Dendritic Cell Identity and Function via Induction of Transcriptional Regulator DC-SCRIPT. Immunity 50, 77–90 e5 (2019).

53. Welner, R.S. et al. C/EBPalpha is required for development of dendritic cell progenitors. Blood 121, 4073–81 (2013).

54. Scholz, F. et al. The transcription factor C/EBPbeta orchestrates dendritic cell maturation and functionality under homeostatic and malignant conditions. Proc Natl Acad Sci U S A 117, 26328–26339 (2020).

55. Hammelman, J., Patel, T., Closser, M., Wichterle, H. & Gifford, D. Ranking reprogramming factors for cell differentiation. Nat Methods 19, 812–822 (2022).

56. Bravo Gonzalez-Blas, C., et al. SCENIC+: single-cell multiomic inference of enhancers and gene regulatory networks. Nat Methods 20, 1355–1367 (2023).

57. Paszke, A. et al. Pytorch: An imperative style, high-performance deep learning library. Advances in neural information processing systems 32(2019).

58. Dao, T. Flashattention-2: Faster attention with better parallelism and work partitioning. arXiv preprint arXiv:2307.08691 (2023).

59. Yang, F. et al. scBERT as a large-scale pretrained deep language model for cell type annotation of single-cell RNA-seq data. Nature Machine Intelligence 4, 852-+ (2022).

60. Lin, T.-Y., Goyal, P., Girshick, R., He, K. & Dollár, P. Focal loss for dense object detection. in Proceedings of the IEEE international conference on computer vision 2980-2988 (2017).

61. Wolf, F.A., Angerer, P. & Theis, F.J. SCANPY: large-scale single-cell gene expression data analysis. Genome biology 19, 1–5 (2018).

62. Wang, G., Sarkar, A., Carbonetto, P. & Stephens, M. A simple new approach to variable selection in regression, with application to genetic fine mapping. J R Stat Soc Series B Stat Methodol 82, 1273–1300 (2020).

63. Noguchi, S. et al. FANTOM5 CAGE profiles of human and mouse samples. Sci Data 4, 170112 (2017).

64. Zhang, X. et al. MYOCD is Required for Cardiomyocyte-like Cells Induction from Human Urine Cells and Fibroblasts Through Remodeling Chromatin. Stem Cell Rev Rep 18, 2414–2430 (2022).

65. Ma, H. et al. The nuclear receptor THRB facilitates differentiation of human PSCs into more mature hepatocytes. Cell Stem Cell 29, 1611 (2022).

66. Pedregosa, F. et al. Scikit-learn: Machine learning in Python. the Journal of machine Learning research 12, 2825–2830 (2011).

67. Sherman, B.T. et al. DAVID: a web server for functional enrichment analysis and functional annotation of gene lists (2021 update). Nucleic Acids Res 50, W216–W221 (2022).

68. Vallot, C. et al. Erosion of X Chromosome Inactivation in Human Pluripotent Cells Initiates with XACT Coating and Depends on a Specific Heterochromatin Landscape. Cell Stem Cell 16, 533–46 (2015).

69. Akdemir, K.C. et al. Genome-wide profiling reveals stimulus-specific functions of p53 during differentiation and DNA damage of human embryonic stem cells. Nucleic Acids Res 42, 205–23 (2014).

70. Rao, S.S. et al. A 3D map of the human genome at kilobase resolution reveals principles of chromatin looping. Cell 159, 1665–80 (2014).

71. Krietenstein, N. et al. Ultrastructural Details of Mammalian Chromosome Architecture. Mol Cell 78, 554–565.e7 (2020).

72. Rusan, M. et al. Suppression of Adaptive Responses to Targeted Cancer Therapy by Transcriptional Repression. Cancer Discov 8, 59–73 (2018).

73. Liu, X. et al. In Situ Capture of Chromatin Interactions by Biotinylated dCas9. Cell 170, 1028–1043.e19 (2017).

74. Maurano, M.T. et al. Large-scale identification of sequence variants influencing human transcription factor occupancy in vivo. Nat Genet 47, 1393–401 (2015).

75. Song, L. et al. Open chromatin defined by DNaseI and FAIRE identifies regulatory elements that shape cell-type identity. Genome Res 21, 1757–67 (2011).

76. Hu, D. et al. The MLL3/MLL4 branches of the COMPASS family function as major histone H3K4 monomethylases at enhancers. Mol Cell Biol 33, 4745–54 (2013).

77. Ferrari, R. et al. Adenovirus small E1A employs the lysine acetylases p300/CBP and tumor suppressor Rb to repress select host genes and promote productive virus infection. Cell Host Microbe 16, 663–76 (2014).

78. Maurano, M.T. et al. Role of DNA Methylation in Modulating Transcription Factor Occupancy. Cell Rep 12, 1184–95 (2015).

79. Bernstein, B.E. et al. The NIH Roadmap Epigenomics Mapping Consortium. Nat Biotechnol 28, 1045–8 (2010).

80. Baranello, L. et al. RNA Polymerase II Regulates Topoisomerase 1 Activity to Favor Efficient Transcription. Cell 165, 357–71 (2016).

81. Singh, Amar M. et al. Cell-Cycle Control of Bivalent Epigenetic Domains Regulates the Exit from Pluripotency. Stem Cell Reports 5, 323–336 (2015).

82. Ma, H. et al. The nuclear receptor THRB facilitates differentiation of human PSCs into more mature hepatocytes. Cell Stem Cell 29, 795–809.e11 (2022).

83. Grant, C.E., Bailey, T.L. & Noble, W.S. FIMO: scanning for occurrences of a given motif. Bioinformatics 27, 1017–8 (2011).

84. Gertz, J. et al. Distinct properties of cell-type-specific and shared transcription factor binding sites. Mol Cell 52, 25–36 (2013).

85. Huang da, W., Sherman, B.T. & Lempicki, R.A. Systematic and integrative analysis of large gene lists using DAVID bioinformatics resources. Nat Protoc 4, 44–57 (2009).

