## Supplementary Figures for "Learning interpretable representation for context-specific transcription regulatory networks using a foundation model"

##### Supplementary Fig. S1

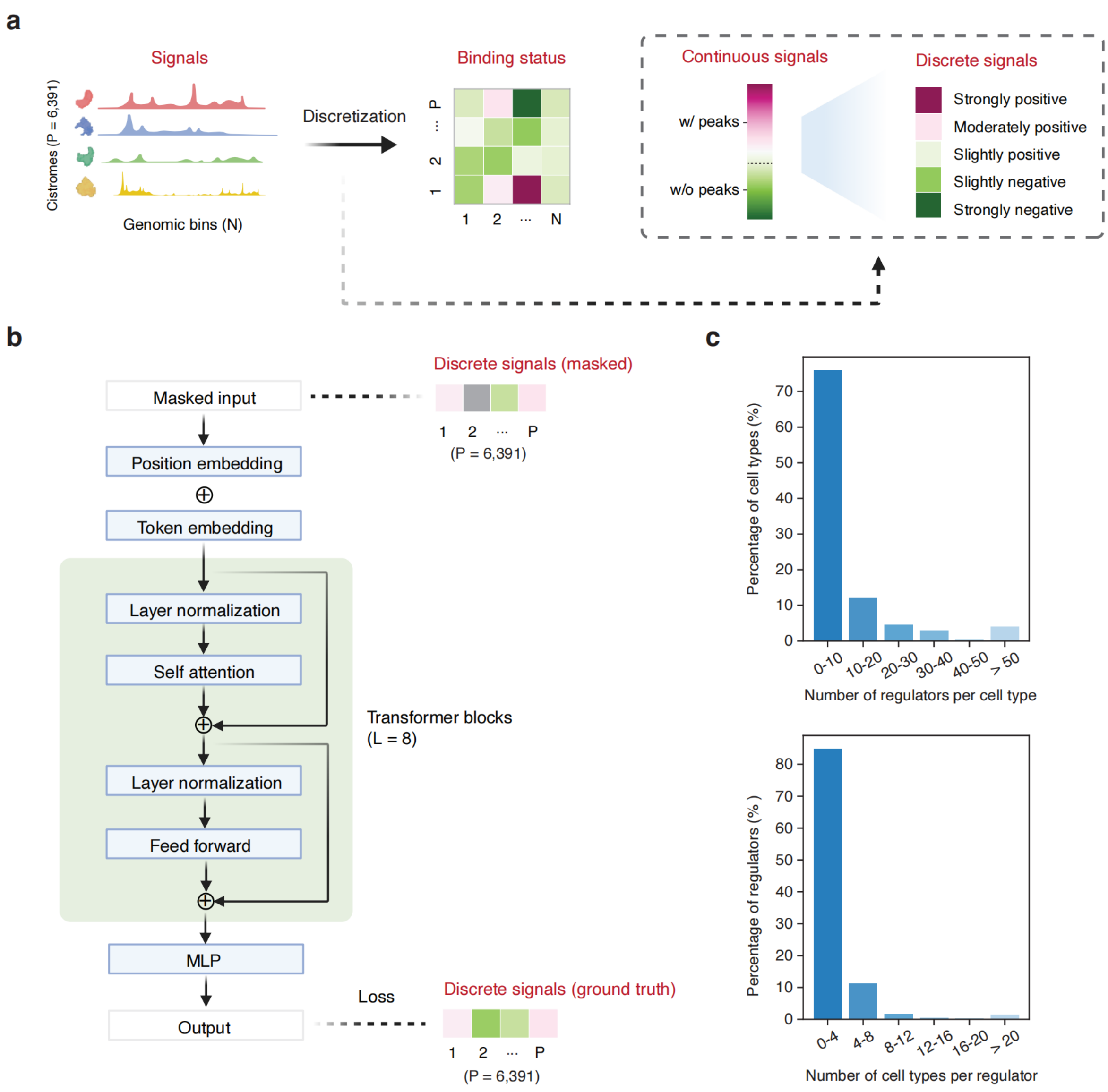


**Supplementary Fig. S1. Pre-training dataset and workflow.**

**a.** Schematic illustrating the tokenization process of the Cistrome-Human-6K datasets. Continuous signals for each cistrome were converted into discrete signals and categorized into five categories from low to high signals (see Methods for details). **b.** Schematic showing pre-trained model architecture of ChromBERT, which includes two embedding layers to handle token (binding status) and positional information (the identity of cistrome), eight transformer blocks designed to capture co-association pattern among cistromes, and an MLP decoder to predict the binding status of cistromes. The model parameters were updated effectively through loss calculations on masked objectives (see Methods for details). **c.** Bar plots showing the data distribution of cell types and regulators in the Cistrome-Human-6K dataset. The majority of cell types have available cistromes for fewer than 10 regulators (top), while most regulators have cistromes present in fewer than 4 cell types (bottom), highlighting the sparsity of the dataset's coverage.

#### Supplementary Fig. S2

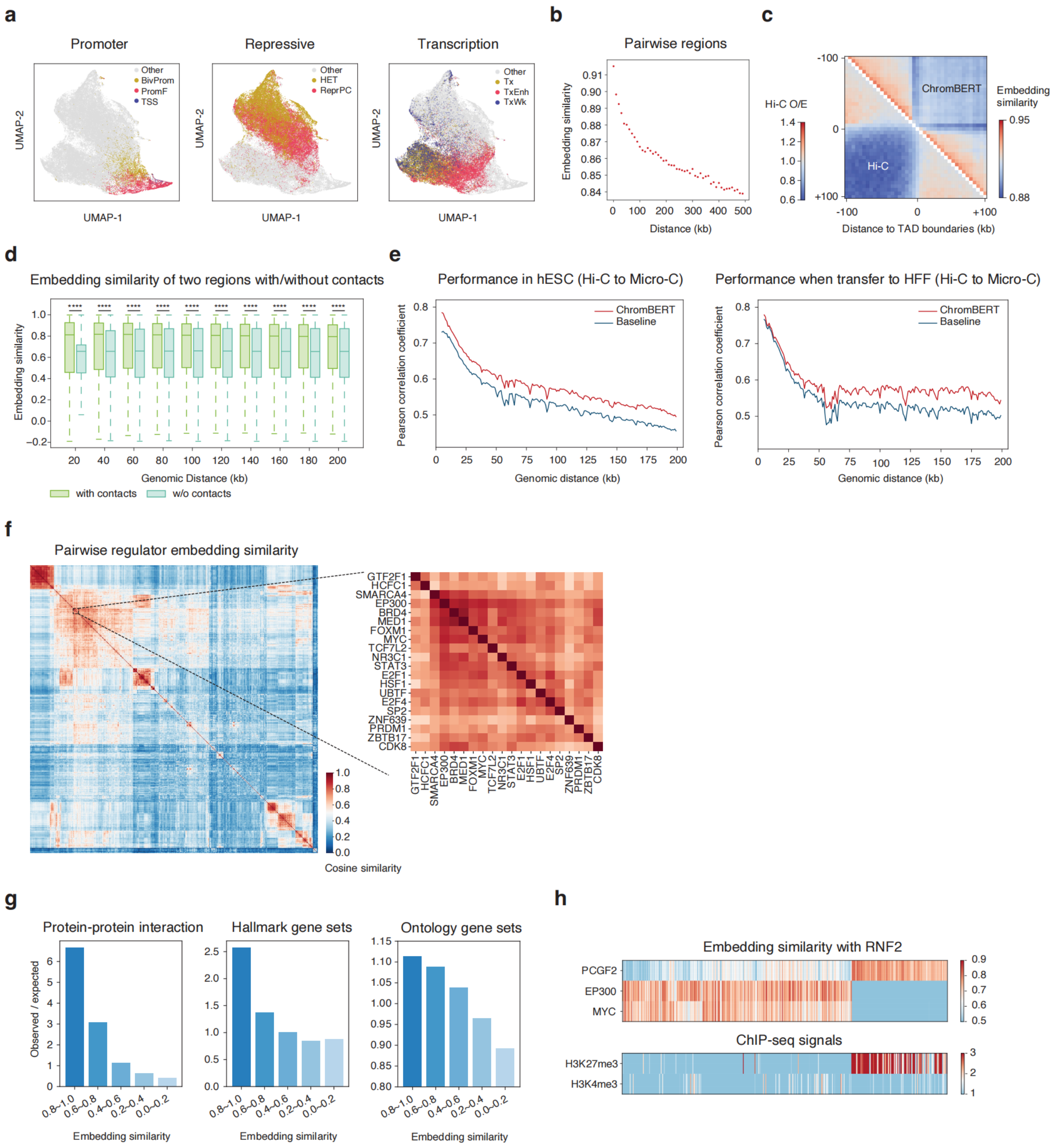


**Supplementary Fig. S2. Pre-trained TRN embeddings and regulator embeddings represent human TRNs.**

**a.** Uniform Manifold Approximation and Projection (UMAP)[1] visualization of ChromBERT's mean pooled TRN embeddings across regions on chromosome one, color-coded by chromatin states as reported in the previous publication[2]. **b.** The averaged pairwise cosine similarity between ChromBERT's aggregated TRN embeddings for 761,838 randomly selected genomic bin pairs from chromosome one, showing a distance decay effect. **c.** Bottom triangle: heatmap showing aggregated Hi-C interactions, indicated by observed/expected (O/E) values, centered around TAD boundaries on chromosome one. Top triangle: heatmap illustrating the pairwise cosine similarity of aggregated TRN embeddings of genomic bins. The used Hi-C data in K562 cells was from the previous study (GSE63525[3]). **d.** Box plots showing the cosine similarity of ChromBERT's mean-pooled TRN embeddings for pairs of genomic regions in hESC cells, stratified by the presence (green) or absence (blue) of Micro-C contacts across different genomic distances. Region pairs with Micro-C contacts consistently exhibit significantly higher cosine similarity than those without at all distances. Statistical significance was assessed using a two-sided Student's *t*-test, with **** indicating *p*-value < $10^{-4}$. The center lines mark the median, the box limits indicate the 25th and 75th percentiles, and the whiskers extend to 1.5× the interquartile range from the 25th and 75th percentiles. **e.** Line plot illustrating the distance-stratified Pearson's correlation coefficient between observed Micro-C contact maps and CNN-predicted contact maps derived from ChromBERT's TRN embeddings and Hi-C contact maps in hESC cells (see Methods for details). The baseline comparison is made using Hi-C contact maps interpolated from 5-kb to 1-kb resolution. The CNN model was trained and tested in hESC cells (left) and directly transferred to HFF cells (right). The Micro-C and Hi-C data used in **d-e** of hESC cells (4DNES21D8SP8, 4DNES2M5JIGV) and HFF cells (4DNESWST3UBH, 4DNES2R6PUEK) were obtained from previous study[4]. **f.** Heatmap depicting the cosine similarities of ChromBERT's regulator embeddings on chromosome one, revealing complex interaction between transcription regulators (left). A zoomed-in view highlights that functionally associated regulators, such as EP300 and BRD4, have higher embedding similarities (right). For each regulator, regulator embeddings were compiled from regions across chromosome one into a 768-dimensional vector, facilitating the comprehensive analysis. **g**. Bar plot shows pre-trained ChromBERT's regulator embeddings can represent the functional collaborations among transcription regulators. Regulator pairs were ranked by the cosine similarity between their averaged embeddings across chromosome one. The observed (each group) / expected (all pairs) ratios for protein-protein interaction frequency (BioPlex[5]), co-occurrence frequency in the same hallmark gene sets or ontology gene sets (Molecular Signatures Database[6]) were shown, highlighting significant functional collaborations among regulators with high embedding similarity. **h**. Top, heatmap shows the embedding similarities between RNF2 and other regulators at representative RNF2 peaks. The color represents the cosine similarity of two regulator embeddings. Bottom, the H3K4me3 and H3K27me3 ChIP-seq profiles at the given genomic regions. The color represents ChIP-seq signals in RPM. RNF2, H3K4me3 and H3K27me3 ChIP-seq data were from previous study (GSE105028[7], GSE39912[8] and GSE62562[9]).

#### Supplementary Fig. S3

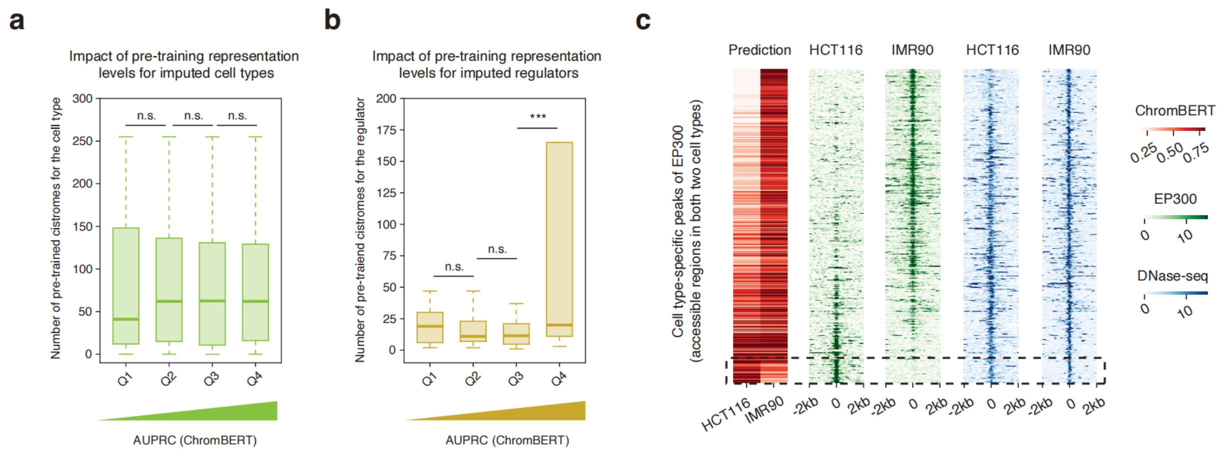


**Supplementary Fig. S3. Robustness of ChromBERT in cistrome imputation.**

**a-b.** The impact of pre-training representation levels on the performance of cistrome imputation. The *X*-axis represents performance quantiles on cistrome imputation, divided into four groups: the first quantile (Q1, 0–25%), second quantile (Q2, 25–50%), third quantile (Q3, 50–75%), and fourth quantile (Q4, 75–100%), ordered from low to high performance. The *Y*-axis indicates the pre-training representation levels (number of cistromes) for the cell type (**a**) or the transcription regulator (**b**) of the test cistrome being imputed. Statistical significance was conducted using a two-sided Student's *t*-test, where n.s. indicates non-significant and *** indicates *p*-value < 0.001. The center lines mark the median, the box limits indicate the 25th and 75th percentiles, and the whiskers extend to 1.5× the interquartile range from the 25th and 75th percentiles. **c.** Heatmaps present ChromBERT's predicted probabilities using DNase-seq prompts (red), alongside ChIP-seq signals (green) and DNase-seq signals (blue) on cell-type-specific peaks of EP300 that were accessible in both HCT116 and IMR90 (see Methods for details). The EP300 ChIP-seq and DNase-seq signals were normalized to the genome average. Black dashed rectangles highlight cell-type-specific binding events that do not differ in DNase-seq signals, yet were accurately predicted by ChromBERT.

#### Supplementary Fig. S4

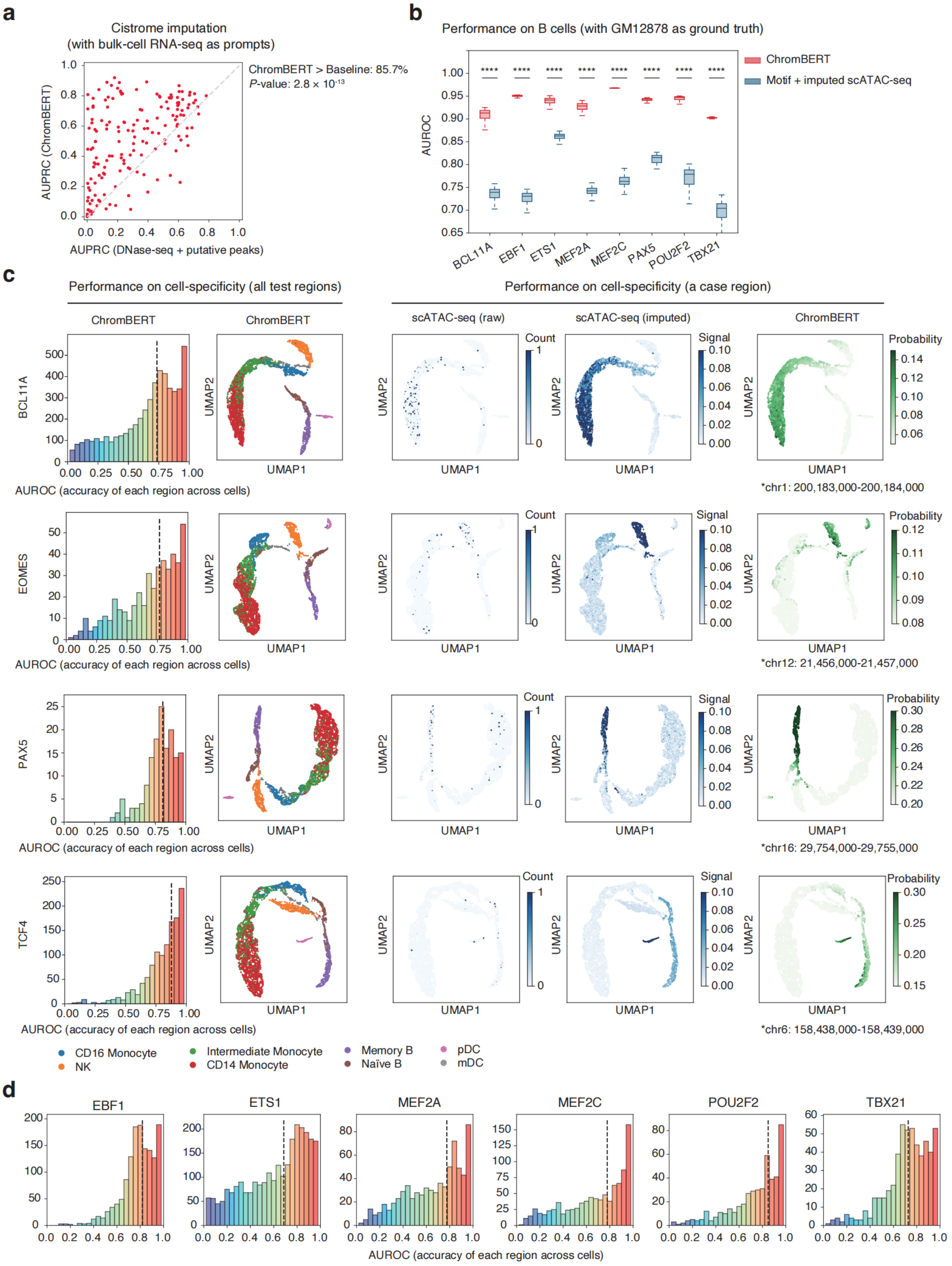

**Supplementary Fig. S4. ChromBERT imputes cistromes at single-cell resolution using scGPT prompts.**

**a**. Scatter plot illustrating the performance of cistrome imputation tasks for ChromBERT using transcriptome prompts in bulk cells, compared to baseline methods measured by DNase-seq signal and putative peaks (see Methods for details). The diagonal gray line indicates equal performance, with percentages of cistromes where ChromBERT outperforms annotated alongside the *p*-value calculated by a two-sided Student's *t*-test. **b.** Box plots comparing prediction performance in B cells using bulk-cell ChIP-seq data from GM12878 as the ground truth. Red boxes represent ChromBERT with single-cell prompts, while blue boxes indicate a combination of DNA-binding motifs and imputed single-cell ATAC-seq signals (via scOpen[10]). The comparison was performed at 1-kb bins containing imputed single-cell ATAC-seq peaks (imputed single-cell ATAC-seq signals > 0.1) in the PBMC dataset and the presence of motifs for each regulator. Statistical significance was computed with the two-sided Student's *t*-test, where **** represents *p*-value < $10^{-4}$. The center lines mark the median, the box limits indicate the 25th and 75th percentiles, and the whiskers extend to 1.5× the interquartile range from the 25th and 75th percentiles. **c**. Cell-specificity analysis of cistrome imputation by ChromBERT with single-cell prompts. The first and second columns evaluate performance across all test regions: the first column shows the distribution of AUROC scores for cell-specificities of each region, and the second column presents a UMAP analysis of predictive probabilities across cells, with cells colored by pseudo-bulk cell type annotations (from paired single-cell RNA-seq). The third to fifth columns focus on the cell-specificity of a case region: the third column shows raw ATAC-seq counts, the fourth displays imputed ATAC-seq counts, and the fifth presents ChromBERT's predictions across different cells. Test regions were defined as a merge set of regions showing high chromatin accessibility specificity in each pseudo-bulk cell type. Each single cell was assigned to a pseudo-bulk cell type based on single-cell RNA-seq annotations. Specific regions for each pseudo-bulk cell type were defined as those accessible (with imputed single-cell ATAC-seq peak) in at least 60% of cells assigned to that cell type and accessible in no more than 10% of other cells, with a mean fold change in accessible cell number between the two groups of cells > 2. All pseudo-bulk cell-type-specific regions were pooled as total test regions for this analysis, while evaluation for each specific regulator was only performed in total regions with the regulator's motif. Pseudo-bulk cell types related to T cells are excluded due to insufficient representation by scGPT. **d**. Cell-specificity analysis of cistrome imputation by ChromBERT with single-cell prompts for additional transcription regulators.

### Supplementary Fig. S5

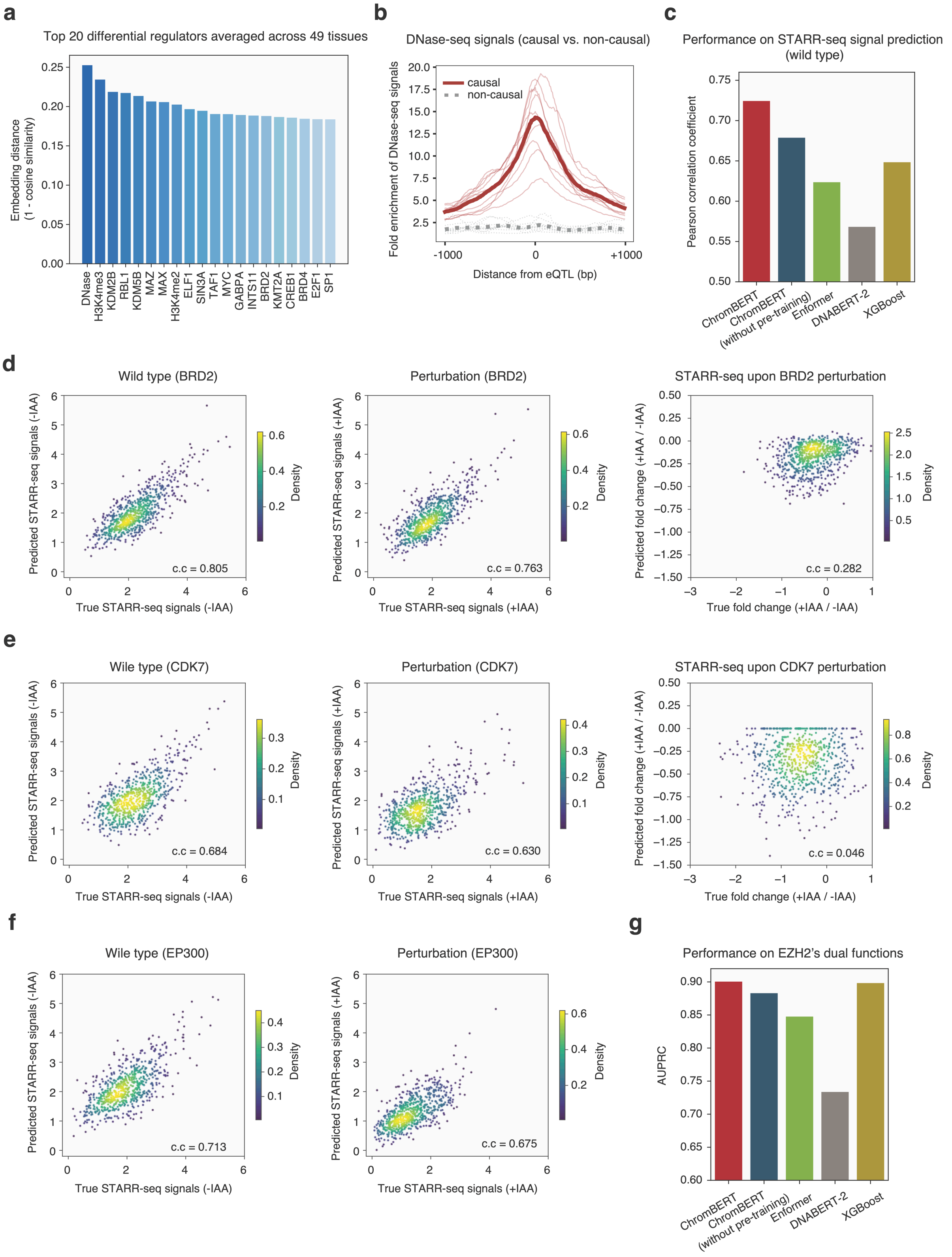


**Supplementary Fig. S5. Context-specificities of TRNs revealed by ChromBERT.**

**a.** Bar plot illustrating the top 20 transcription regulators ranked by distance in regulator embeddings between causal and non-causal eQTLs, averaged across 49 tissues. **b.** Average DNase-seq site profiles around causal versus non-causal eQTLs. Each thin line represents an individual tissue (10 tissues used, see Data Availability for details), while the bold line represents the mean profiles across 10 tissues. Red lines indicate causal eQTLs and gray dashed lines indicate non-causal eQTLs. **c.** Bar plots illustrate the performance of ChromBERT, ChromBERT (without pre-training), Enformer, DNABERT-2, and XGBoost in modeling STARR-seq signals in wild type HCT116 cells. **d-f.** Scatter plots showing the prediction performance of ChromBERT for in silico perturbation studies. The predicted and ground truth wild type STARR-seq signals (left), IAA-treated (perturbation of factors) STARR-seq signals (center) and $log_2$-transformed fold change of perturbation versus wild-type (right) were shown, respectively. Analyses were conducted using STARR-seq datasets with perturbation for three cofactors, BRD2 (**d**), CDK7 (**e**), and EP300/CREBBP (**f**) (GSE156740[11]). Pearson's correlation coefficients were annotated in the plot, and the color represents the density of points. **g.** Bar plots show the performance of ChromBERT, ChromBERT (without pre-training), Enformer, DNABERT-2 and XGBoost in classifying EZH2's classical and non-classical sites. ChromBERT was fine-tuned by omitting cistromes associated with H3K27me3 from input reference cistromes to diminish its dominant role in the task. EZH2 and H3K27me3 ChIP-seq data were from previous studies (GSE61176 and GSE29611[12]).

#### Supplementary Fig. S6

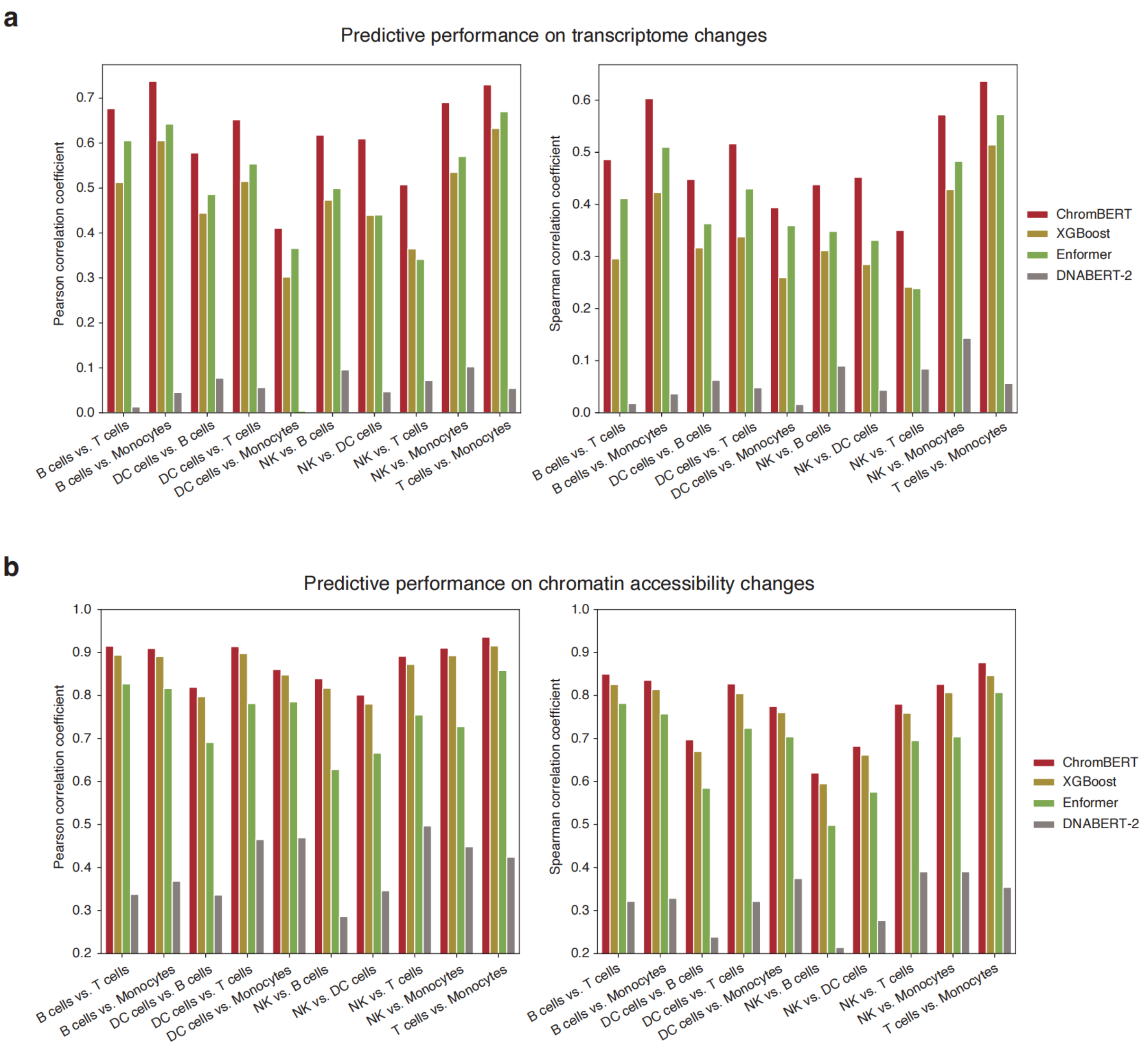


**Supplementary Fig. S6. Benchmark on prediction for transcriptome and chromatin accessibility changes between single-cell pseudo-bulk cell types.**

**a, b.** Bar plots comparing performances of ChromBERT, XGBoost, Enformer and DNABERT-2 in predictions for genome-wide transcriptome (**a**) and chromatin accessibility (**b**) changes between different single-cell pseudo-bulk cell types. Pearson's correlation coefficient and Spearman's correlation coefficient were computed and shown.

#### Supplementary Fig. S7

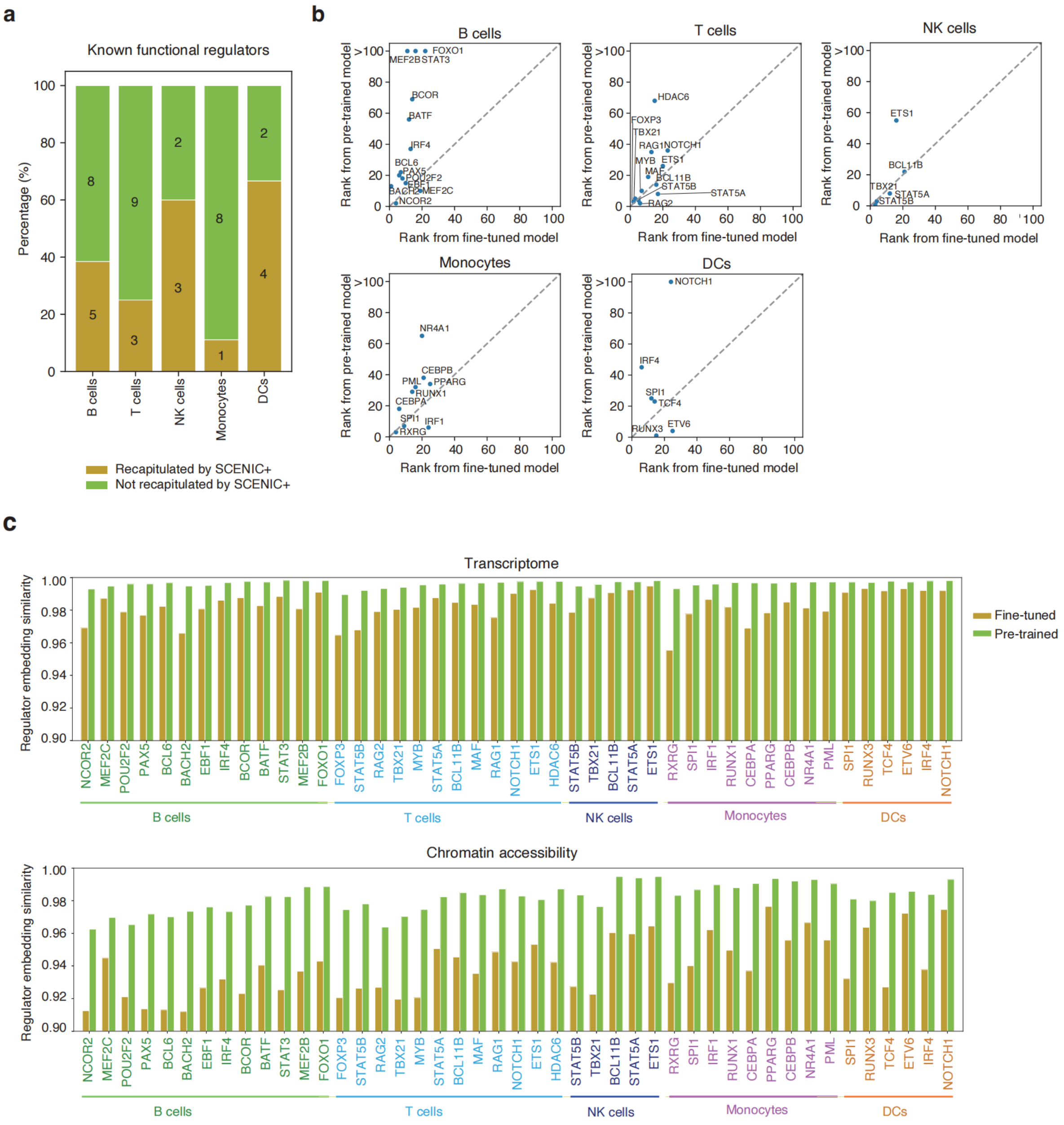


**Supplementary Fig. S7. Identified key regulators for cellular heterogeneity.**

**a.** Bar plots illustrating the percentage of known key regulators identified by ChromBERT in various cell types that were not detected by SCENIC+ (v1.0.1)[13]. **b.** Scatter plots comparing the rankings of known key regulators using fine-tuned regulator embeddings (*X*-axis) versus pre-trained regulator embeddings (*Y*-axis). **c.** Bar plots showing the comparison of embedding similarity for known key regulators between regions with increased activity and unchanged activity. The plots reveal that known key regulators exhibit lower embedding similarity with fine-tuned regulator embeddings compared to pre-trained regulator embeddings, suggesting that fine-tune model can capture important regulator in determining cellular heterogeneity.

#### Supplementary Fig. S8

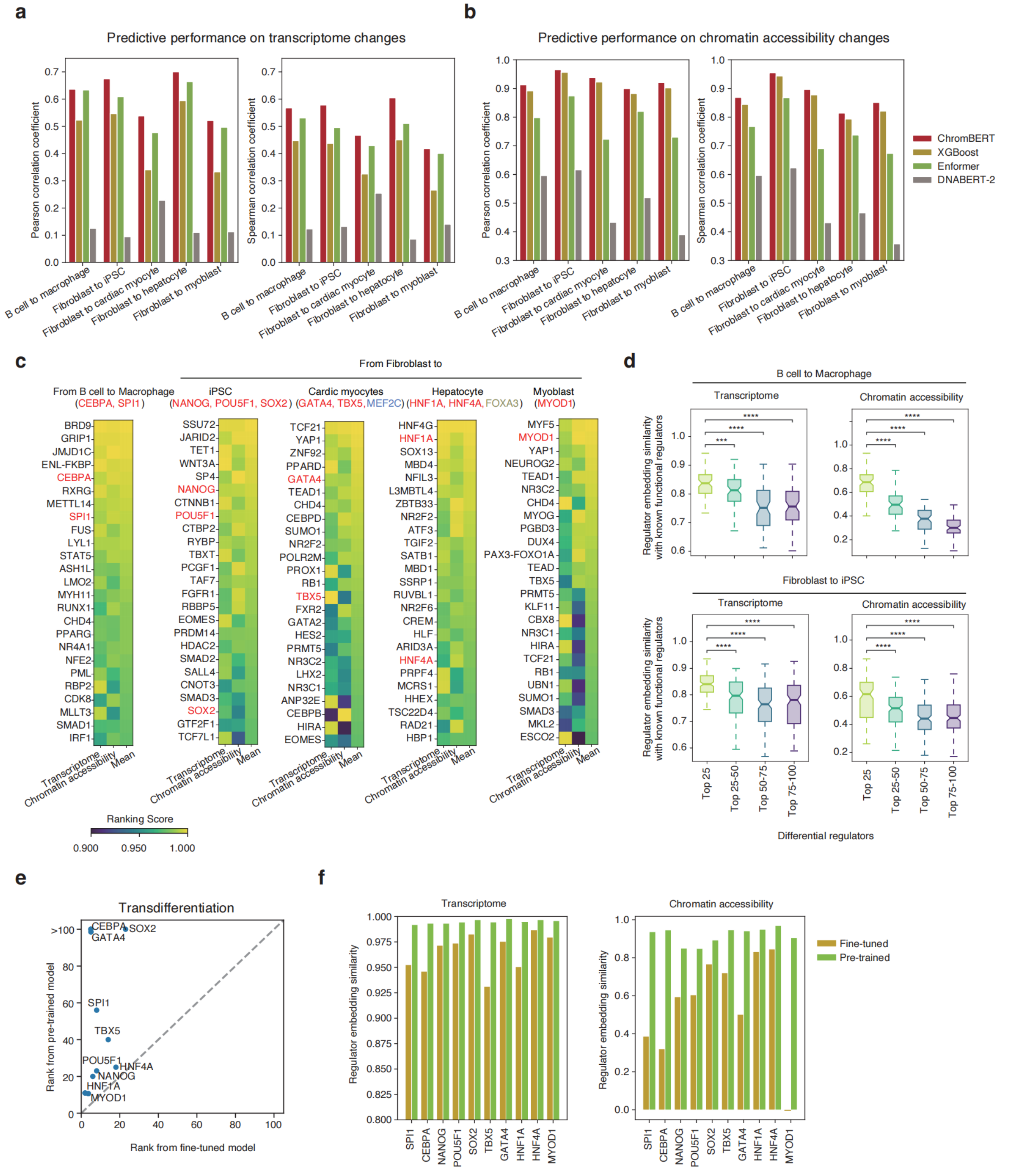


**Supplementary Fig. S8. Identified key regulators for transdifferentiation.**

**a, b.** Bar plots comparing performances of ChromBERT, XGboost, Enformer and DNABERT-2 in predictions for genome-wide transcriptome (**a**) and chromatin accessibility (**b**) changes between different cell states. Pearson's correlation coefficient and Spearman's correlation coefficient were computed and shown. **c.** Heatmap shows ranking scores of the top 25 regulators identified by ChromBERT. The known driver regulators of each transdifferentiation process were annotated above the heatmaps and known drivers successfully recapitulated by ChromBERT were highlighted in red. **d.** Box plots showing embedding similarity between top-ranked regulators (w/o known driver regulators) and known driver regulators, suggesting the top-ranked regulators have high potential function associations with known driver regulators. Significance tests were performed by using two-sided Student's *t*-tests. *** represents *p*-value < 0.001, **** represents *p*-value < $1 \times 10^{-4}$ and ns represents non-significant. **e.** Scatter plots comparing the ranking of known driver regulators by interpreting fine-tuned regulator embeddings (*X*-axis) or pretrained regulator embeddings (*Y*-axis) (see Methods for details). **f.** Bar plots comparing the embedding similarity of known driver regulators between regions with increased activity and unchanged activity, highlighting that known driver regulators show lower embedding similarity with fine-tuned regulator embeddings than pre-trained regulator embeddings. It suggests that fine-tuned model can capture important regulators in determining cellular state transition more effectively.

#### Supplementary Tables

Supplementary Table S1. Metadata for the Cistrome-Human-6K dataset.

Supplementary Table S2. Performance on cistrome imputation in unseen cell types.

Supplementary Table S3. Hyperparameters used for fine-tuning.

Supplementary Table S4. Metadata for training the prompt-enhanced ChromBERT of cistrome imputation.

#### Supplementary References


1. Becht, E. *et al.* Dimensionality reduction for visualizing single-cell data using UMAP. *Nat Biotechnol* (2018).
2. Vu, H. & Ernst, J. Universal annotation of the human genome through integration of over a thousand epigenomic datasets. *Genome Biol* **23**, 9 (2022).
3. Rao, S.S. *et al.* A 3D map of the human genome at kilobase resolution reveals principles of chromatin looping. *Cell* **159**, 1665-80 (2014).
4. Krietenstein, N. *et al.* Ultrastructural Details of Mammalian Chromosome Architecture. *Mol Cell* **78**, 554-565.e7 (2020).
5. Huttlin, E.L. *et al.* Dual proteome-scale networks reveal cell-specific remodeling of the human interactome. *Cell* **184**, 3022-3040 e28 (2021).
6. Liberzon, A. *et al.* The Molecular Signatures Database (MSigDB) hallmark gene set collection. *Cell Syst* **1**, 417-425 (2015).
7. Lyu, X., Rowley, M.J. & Corces, V.G. Architectural Proteins and Pluripotency Factors Cooperate to Orchestrate the Transcriptional Response of hESCs to Temperature Stress. *Mol Cell* **71**, 940-955 e7 (2018).
8. Vallot, C. *et al.* Erosion of X Chromosome Inactivation in Human Pluripotent Cells Initiates with XACT Coating and Depends on a Specific Heterochromatin Landscape. *Cell Stem Cell* **16**, 533-46 (2015).
9. Akdemir, K.C. *et al.* Genome-wide profiling reveals stimulus-specific functions of p53 during differentiation and DNA damage of human embryonic stem cells. *Nucleic Acids Res* **42**, 205-23 (2014).
10. Li, Z. *et al.* Chromatin-accessibility estimation from single-cell ATAC-seq data with scOpen. *Nat Commun* **12**, 6386 (2021).
11. Neumayr, C. *et al.* Differential cofactor dependencies define distinct types of human enhancers. *Nature* **606**, 406-413 (2022).

12. Consortium, E.P. *et al.* Expanded encyclopaedias of DNA elements in the human and mouse genomes. *Nature* **583**, 699-710 (2020).
13. Bravo Gonzalez-Blas, C. *et al.* SCENIC+: single-cell multiomic inference of enhancers and gene regulatory networks. *Nat Methods* **20**, 1355-1367 (2023).
